# A systems approach reveals species differences in hepatic stress response capacity

**DOI:** 10.1101/2022.04.13.488145

**Authors:** Giusy Russomanno, Rowena Sison-Young, Lucia A. Livoti, Hannah Coghlan, Rosalind E. Jenkins, Steven J. Kunnen, Ciarán P. Fisher, Dennis Reddyhoff, Iain Gardner, Adeeb H. Rehman, Stephen W. Fenwick, Andrew R. Jones, Guy Vermeil De Conchard, Gilles Simonin, Helene Bertheux, Richard J. Weaver, Michael J. Liguori, Diana Clausznitzer, James L. Stevens, Christopher E. Goldring, Ian M. Copple

## Abstract

To minimise unexpected toxicities in early phase clinical studies of new drugs, it is vital to understand fundamental similarities and differences between preclinical test species and humans. We have used physiologically-based pharmacokinetic modelling to identify doses of the model hepatotoxin acetaminophen yielding similar hepatic burdens of the reactive metabolite N-acetyl-p-benzoquinoneimine in mice and rats, to enable comparison of tissue adaptive responses under conditions of equivalent chemical insult. Mice exhibited a greater degree of liver injury than rats, despite the equivalent hepatic NAPQI burden. Transcriptomic and proteomic analyses highlighted the stronger activation of stress response pathways (including the Nrf2 oxidative stress response and autophagy) in the livers of rats. Components of these pathways were also found to be expressed at a higher basal level in the livers of rats compared with both mice and humans. Our findings exemplify a systems approach to understanding differential species sensitivity to hepatotoxicity, and have important implications for species selection and human translation in the safety testing of new drug candidates.

## INTRODUCTION

A critical stage in the development of a new medicine is the transition from preclinical experiments to trials in humans. The success of this transition is dependent on the robust translation of preclinical findings to clinical responses, in terms of both efficacy and safety ^1,2,3^. For example, there are numerous cases where a drug candidate has caused serious liver toxicity in patients despite preclinical safety assessment indicating no cause for concern ^4,5,6^. On the other hand, many developmental compounds deemed unsafe in preclinical species may have been found to be safe in humans. In order to minimise the occurrence of unexpected clinical toxicities and improve the efficiency of the drug development process, is it critical that decision making is informed by a high degree of confidence in preclinical safety data. Related to this, it is vital to understand fundamental similarities and differences amongst and between preclinical species and humans, to inform the selection of appropriate preclinical models, and so accurately predict clinical responses based on in vivo and in vitro data. Conceptually, differential adverse effects of a drug in different species may be a result of dissimilar toxicokinetics and/or toxicodynamics. More specifically, the differential response to a drug may be driven by (a) differences in its disposition (e.g. a toxic metabolite is formed to a greater extent in the sensitive species, such as with the hepatotoxin acetaminophen (APAP ^7^), (b) the differential expression of a target protein (e.g. the equilibrative nucleoside transporter 1, in the case of fialuridine liver toxicity ^8^) or (c) fundamental differences in the nature and/or extent of the downstream cellular response to chemical insult. Whilst (a) and (b) have been extensively studied in the context of various drug toxicities, (c) has garnered less attention to date.

APAP is the single largest cause of acute liver failure in the UK and US ^6, 9^, and is commonly used as an exemplar drug for studying hepatotoxicity. APAP is metabolically bioactivated to the highly reactive N-acetyl-p-benzoquinonimine (NAPQI) which depletes glutathione (GSH) stores, induces oxidative stress and covalently reacts with hepatocellular proteins leading to necrosis ^10, 11^. It is well known that mice (oral LD50 ∼350 mg/kg) are relatively sensitive to APAP liver injury compared with rats (oral LD50 ∼2000 mg/kg) ^7^. Previous work has indicated that this disparity is related to species differences in metabolism, with mice exhibiting a higher fraction of APAP bioactivated to NAPQI ^12,13,14^. To account for this species difference in metabolism, and better design a comparative study of the response of mice and rats to NAPQI insult, we have used physiologically-based pharmacokinetic (PBPK) modelling and simulation to identify doses of APAP yielding an equivalent hepatic burden of NAPQI in the two species. We combined this with a systems approach to compare changes in the hepatic transcriptome and proteome of mice and rats following exposure to equivalent NAPQI burden, and reveal marked differences between the rodent species, and humans, in the basal and adaptive capacities of hepatic stress response pathways that are known to influence APAP toxicity and other forms of drug-induced liver injury (DILI). Our findings have important implications for the selection of preclinical models during the toxicity assessment of new drug candidates.

## RESULTS

### Identification of pharmacokinetically-equivalent doses of APAP in mice and rats

Although modelling the impact on drug exposure of species differences in metabolism is routine, to date little has been done to rigorously compare tissue responses to a given chemical entity across preclinical species in order to identify the species that can better translate to humans. Our preliminary 24 h dose-ranging study with the model hepatotoxin APAP in male C57Bl/6J mice and Sprague-Dawley rats confirmed previous reports ^7^ that equivalent liver injury could not be achieved in the two species, on the basis of changes in serum biomarkers and the extent of centrilobular hepatocellular degeneration/necrosis (Supplementary Fig. 1 and 2). This study, in agreement with previous reports ^10^, confirmed that 300 mg/kg is a non-lethal, hepatotoxic dose of APAP in mice (Supplementary Fig. 1 and 2). We therefore used PBPK modelling and simulation to identify 300 and 1000 mg/kg APAP as oral equivalent doses (OEDs) resulting in an equivalent hepatic NAPQI burden in mice and rats, respectively (Fig. 1). We used these doses of APAP to investigate the temporal liver tissue responses of the two species under conditions of equivalent NAPQI insult. In mice, 300 mg/kg APAP resulted in significant elevations of serum ALT, AST activities and TBIL concentration that peaked 3 h after dosing (Fig. 2a-c). The response to oral administration of APAP was more rapid than previous reports ^15, 16^, and likely related to enhanced absorption associated with the 1 % hydroxyethylcellulose vehicle. Whilst significant elevations of serum ALT, AST activities and TBIL concentration were also observed in rats dosed with 1000 mg/kg APAP, the peak of the response was detected 9 h after dosing (i.e. later than in mice), and the magnitude of change relative to vehicle treated controls was muted by comparison (Fig. 2a-c). Microscopic findings correlated with these serum biomarker changes. In mice, acute degeneration of centrilobular hepatocytes was observed as early as 2 h post-dose (Fig. 2d-e, Supplementary Fig. 3 and 4). Degeneration progressed to hepatocellular necrosis over time with peak severity observed at 9 h post-dose. In rats, centrilobular hepatocyte necrosis was only observed at 9 to 24 h post-dose and occurred at lower incidence and severity relative to this finding in mice (Fig. 2d-e, Supplementary Fig. 3 and 4). As demonstrated by serum ALT and AST activities returning toward baseline, APAP liver injury was resolving by 24 h in rats, but remained relatively higher at the same time point in mice (Fig. 2a-e). Taken together, these findings demonstrate that hepatocellular injury occurs more rapidly and to a greater extent in mice relative to rats when challenged with an equivalent hepatic NAPQI burden.

**Figure 1.**
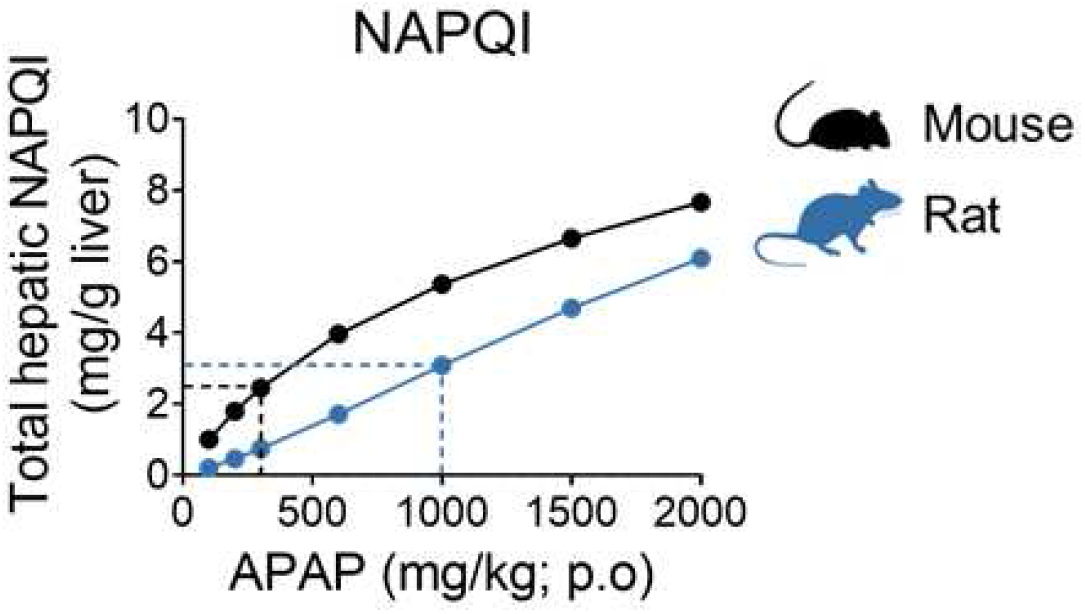
Identification of pharmacokinetically equivalent doses of APAP in mice and rats. Oral equivalent doses (OEDs) of APAP derived from PBPK model simulations that were predicted to give similar levels of total hepatic NAPQI burden in mice (300 mg/kg APAP) and rats (1000 mg/kg APAP).

**Figure 2.**
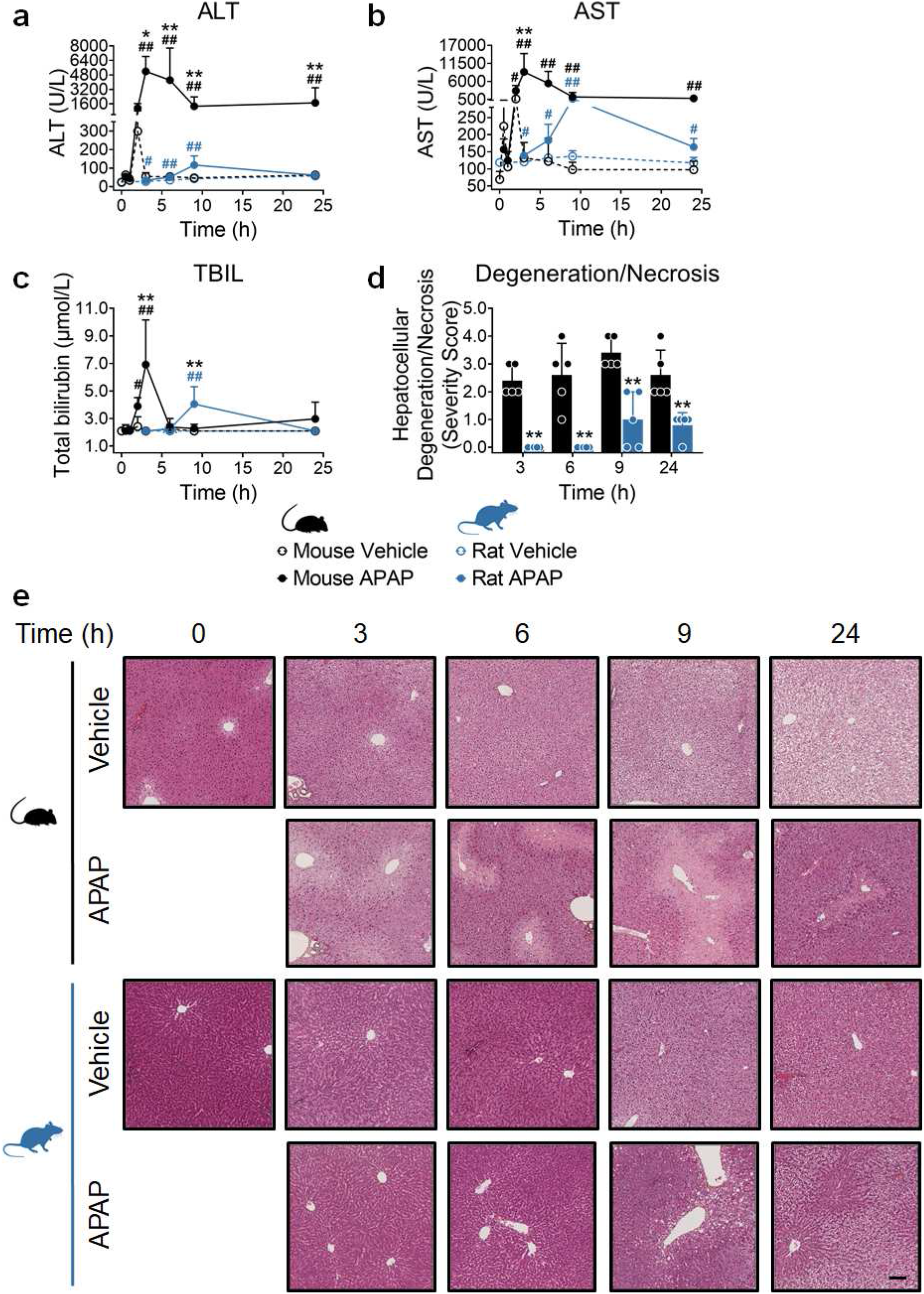
Hepatocellular injury after equivalent hepatic NAPQI burden occurs more quickly and to a greater magnitude in mice compared to rats. Liver injury serum markers (ALT (**a**), AST (**b**) and total bilirubin (TBIL, **c**)) and hepatocellular degeneration/necrosis (**d**) in C57Bl/6J mice (0.5, 1, 2, 3, 6, 9 and 24 h) and Sprague-Dawley rats (3, 6, 9 and 24 h) exposed to 300 mg/kg and 1000 mg/kg APAP, respectively. Time-matched vehicle control groups (dotted lines) were given 1 % (w/v) hydroxyethylcellulose. The extent of centrilobular hepatocellular degeneration/necrosis was assigned a semi-quantitative score as follows: 0 = none observed, 1 = minimal, 2 = mild, 3 = moderate, 4 = marked and 5 = severe (see methods for full details). Values are mean ± SD (n=5). In (**d**) severity scores are shown for each animal. P-values are denoted as ^#^p<0.05, and ^##^p<0.01, comparison of APAP-treated against time-matched control animals (black for mice, blue for rats), or *p<0.05, and **p<0.01, comparison of APAP-treated mice versus rats (Mann-Whitney U test). (**e**) Representative images of haematin-eosin saffron stained liver sections from mice and rats at baseline (0 h) or treated with vehicle or APAP at the indicated time points. Scale bar = 100 μm. See Supplementary Fig. 3 and 4 for more details.

### Confirmation of equivalent chemical insult in the livers of mice and rats

NAPQI is inherently unstable and difficult to quantify bioanalytically. Therefore, as NAPQI is known to deplete hepatic GSH as a prerequisite for liver injury ^10, 11^, we first quantified levels of hepatic GSH in order to confirm the equivalence of chemical insult at the doses of APAP used in mice and rats. In both species, there was a time-dependent increase in GSH in vehicle-treated animals (Fig. 3a), reflecting the restoration of normal resting levels following the reintroduction of food after the 16 h period of fasting prior to APAP dosing. Consistent with the metabolic bioactivation of APAP to NAPQI, there was a significant and equivalent early reduction in GSH in mice and rats treated with APAP (Fig. 3a). In mice, this early depletion was followed by a time-dependent increase in GSH, which reached 55 % of the level quantified in the livers of vehicle-treated controls at 24 h (Fig. 3a). In rats, however, GSH was restored to 86 % of the level of vehicle controls by this point (Fig. 3a), indicating an enhanced capacity of the rat to adapt to the equivalent chemical insult. In addition, pharmacokinetic analysis of the plasma levels of the GSH, N-acetylcysteine and cysteine conjugates of APAP revealed similar levels in the two species (Supplementary Fig. 5). In keeping with the larger dose of APAP administered to rats, both the glucuronide and sulfate conjugates were detected at higher concentrations in time-matched plasma samples compared with mice (Supplementary Fig. 5).

**Figure 3.**
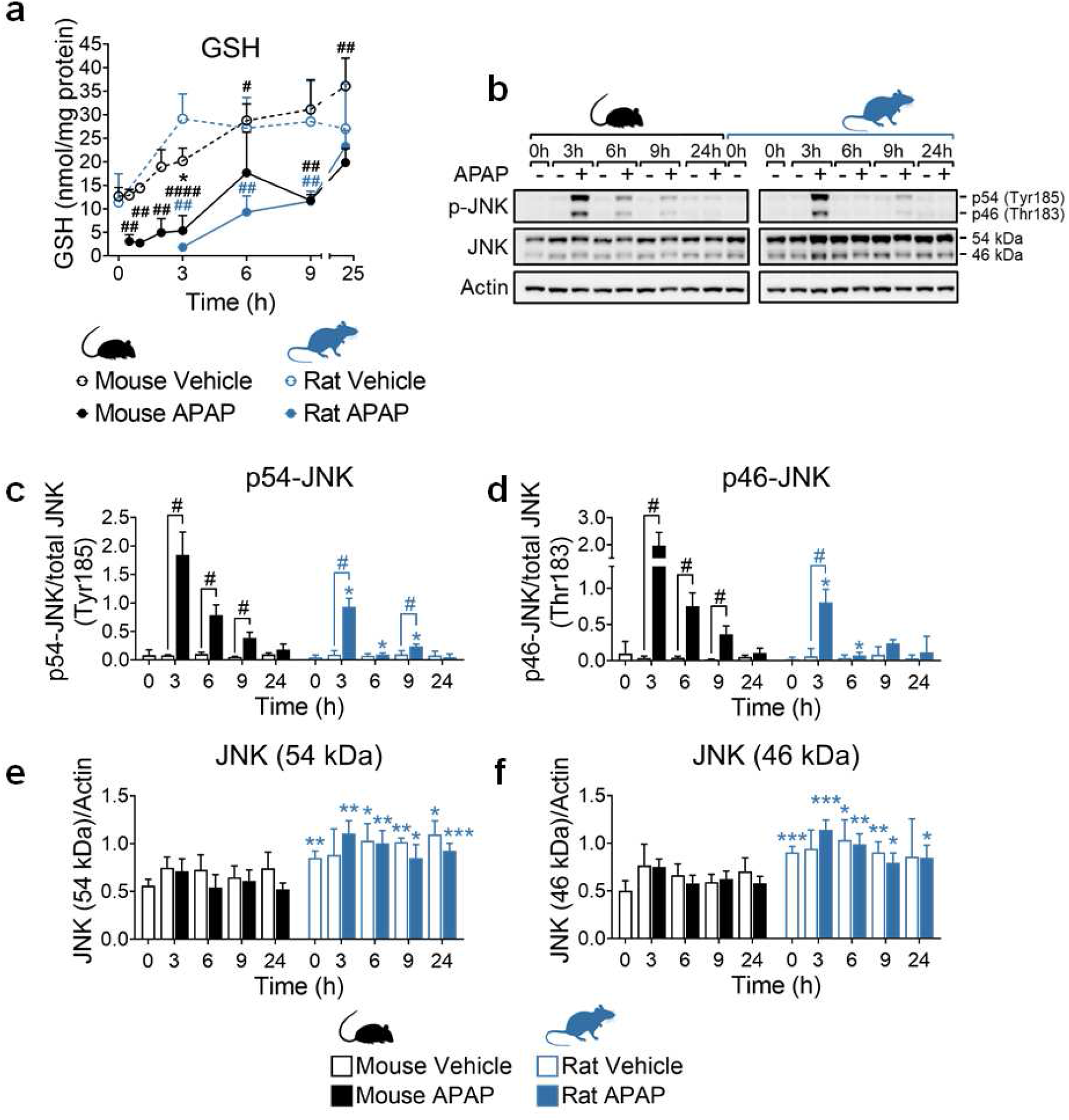
Confirmation of equivalent chemical insult in mice and rats. (**a**) Total hepatic GSH in C57Bl/6J mice (0.5, 1, 2, 3, 6, 9 and 24 h) and Sprague-Dawley rats (3, 6, 9 and 24 h) exposed to 300 mg/kg and 1000 mg/kg APAP respectively. Values are mean ± SD (n=5-12). (**b**) APAP-induced phosphorylation of c-Jun N-terminal kinase (JNK) at Tyr185 (p56) and Thr183 (p46). Samples are pooled from 5 animals per time point. (**c-f**) Densitometric analysis of immunoblots. Protein levels were normalised to total JNK (**c-d**) or β-actin (**e-f**). Values are mean ± SD (n=4). Unpaired t-test or Mann-Whitney U test, as appropriate. P-values are denoted as *p<0.05, **p<0.01, and ***p<0.001, comparison of mouse versus rat for each condition/time point, or ^#^p<0.05, ^##^p<0.01, and ^####^p<0.0001, comparison of APAP-treated animals versus vehicle controls.

Total hepatic NAPQI burden was calculated to assess whether PBPK modelling and simulation had indeed predicted OEDs between the two species. Based on the extent of GSH depletion, and assuming a 1:1 stoichiometry between NAPQI and GSH, total hepatic NAPQI burdens of 1.15 and 0.87 mg/g liver were calculated for mouse and rat, respectively (Supplementary File 1). These calculations from the study data indicate that the PBPK-predicted OEDs for mouse and rat, 300 mg/kg and 1000 mg/kg, respectively, achieved total hepatic NAPQI burdens within 1.3-fold between the two species. Absolute PBPK predictions of total hepatic NAPQI burden were within 3.6-fold of values derived from experimental data for both species (predicted vs. experimental total hepatic NAPQI; 3.19 vs. 1.50 mg for mouse, 27.73 vs. 7.83 mg for rat).

In a further demonstration of the comparable extent of chemical insult in the two species, we detected phosphorylation of c-Jun N-terminal kinase (p-JNK) in mice and rats peaking 3 h after APAP administration (Fig. 3b-d). Phosphorylation and subsequent mitochondrial translocation of JNK is reported to be a critical step in the mechanism of APAP hepatotoxicity ^7^. Whilst the levels of p54- and p46-JNK appeared relatively higher in mice when normalised to the total JNK level, this was largely influenced by the higher basal level of total JNK in rats (Fig. 3b, e-f), as reported by others ^7^. Together, these data confirm the equivalence of chemical insult in the livers of mice and rats treated with 300 and 1000 mg/kg of APAP, respectively.

### Rats exhibit a more robust activation of adaptive stress response pathways than mice in response to an equivalent NAPQI insult

To explore the tissue responses of mice and rats to an equivalent NAPQI insult, we performed transcriptomic analysis on liver tissue collected from each species following administration of vehicle or the OEDs of APAP. The expression of several hundred genes was commonly altered in both mice and rats over the 24 h study period (Fig. 4a), indicating a degree of similarity in the transcriptional response to APAP across the species. However, whilst approximately 300 genes were uniquely responsive to APAP at each time point in the mouse, a far greater number of genes (peaking at 1819 genes by 9 h post-APAP administration) were found to be differentially expressed only in the rat (Fig. 4a), suggesting a more robust overall transcriptional response to the equivalent NAPQI insult in the latter species. Consistent with the relative degrees of drug-induced tissue injury observed in the two species, mice were found to exhibit a stronger early activation of processes involved in regulation of cell death, apoptosis and inflammatory and immune responses, compared with rats (Fig. 4b-d). Moreover, the activation of these processes and associated genes had diminished by 24 h after APAP administration in rats but not in mice (Supplementary Fig. 6), suggestive of a more robust adaptation in the former species. On the other hand, rats exhibited a stronger activation of the Nrf2-driven response to chemical and oxidative stress, along with markers of the endoplasmic reticulum stress response and autophagy, which is a catabolic process that enables the recycling of cellular components and damaged organelles under conditions of stress ^17^ (Fig. 4b-d, Supplementary Fig. 6, Supplementary File 2). The altered expression of representative genes from these processes was confirmed at each time point in the two species via qPCR (Supplementary Fig. 7), which showed an excellent correlation with the corresponding transcriptomics data (Supplementary Fig. 8). We also confirmed that the basal expression level of Nrf2-regulated genes was not differentially influenced by the heavy metal content of the rodent diet ^18^ (Supplementary Fig. 9), in light of previous reports that arsenic can modulate Nrf2 activity ^19, 20^. Our findings were further confirmed by weighted gene co-expression network analysis (WGCNA) using the liver TXG-MAPr tool (https://txg-mapr.eu/, manuscript in preparation). Rats showed a higher average absolute eigengene score (EGs) across all modules (Fig. 5a) and, whilst the overall module responses were well correlated between the two species, responses were typically stronger in rats compared with mice (Fig. 5b). Indeed, rats exhibited a higher perturbation of modules associated with oxidative and endoplasmic reticulum stress responses, as well as autophagy and other adaptive processes (e.g. ribosome biogenesis, which is associated with hepatocellular hypertrophy ^21^) when compared to mice (Fig. 5b-c, Supplementary Fig. 10, Supplementary File 3). Together, these data indicate that the more robust activation of Nrf2 and other adaptive stress responses in rats reflects a fundamental species difference in the tissue response to the equivalent NAPQI insult.

**Figure 4.**
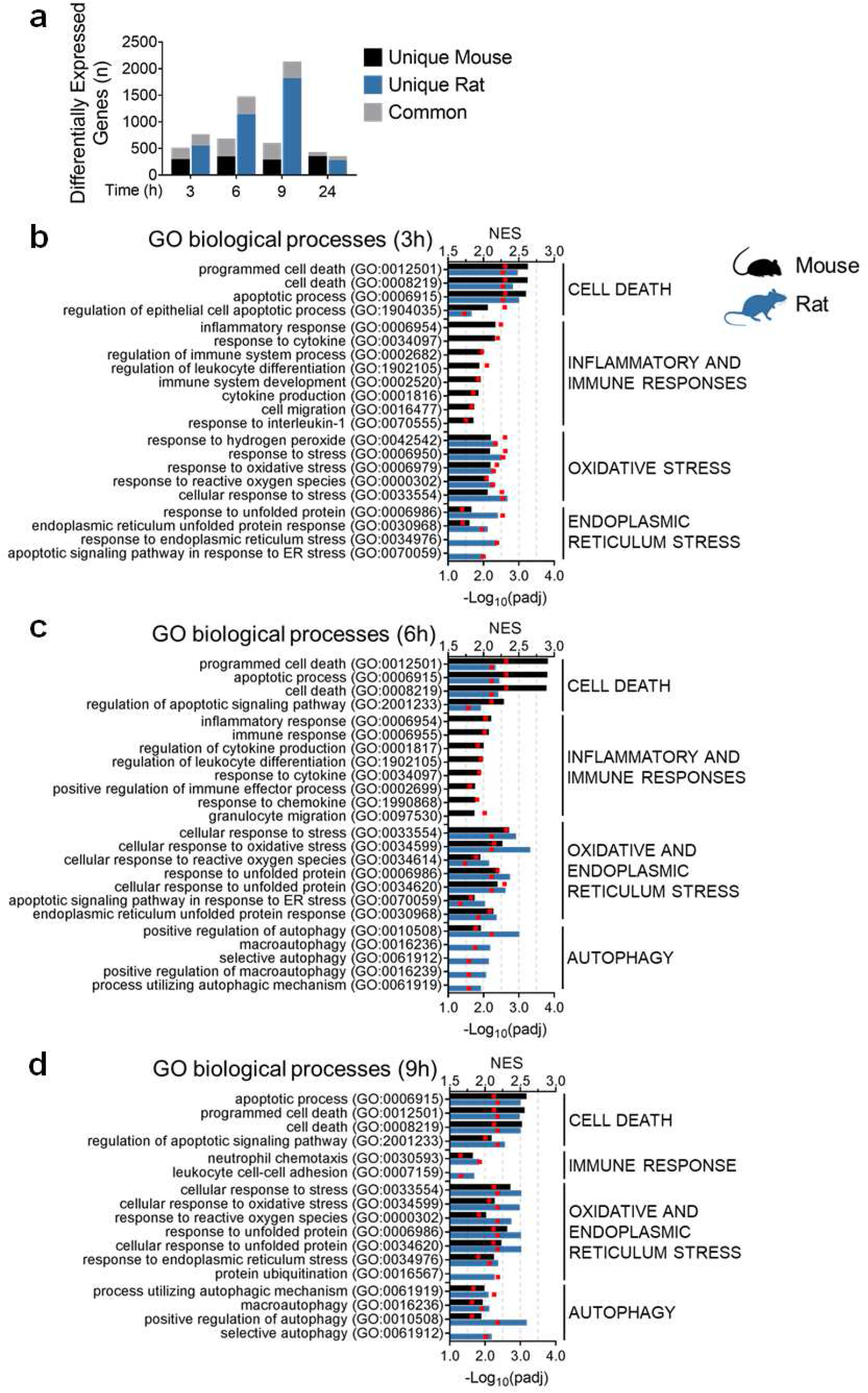
Transcriptomic analysis reveals a more robust activation of adaptive stress response pathways in the liver of APAP-treated rats, compared with mice. (**a**) Number of Differentially Expressed Genes (DEGs; P_adj_ < 0.01, fold change > 1.5 or < −1.5; comparison against time matched vehicle controls). Black and blue bars represent the number of genes that were uniquely responsive to APAP in mice and rat, respectively. Stacked grey bars indicate DEGs in common between the two species. Gene Set Enrichment Analysis (GSEA) at (**b**) 3, (**c**) 6, and (**d**) 9 hours after APAP treatment. Bars represent normalised enrichment scores (NES) of selected gene onotology (GO) terms for both species (mouse in black, rat in blue). Individual GO terms (P_adj_ < 0.05) were clustered into parent terms. The bar is missing for non-significant GO terms. Red squares represent −log_10_ of P_adj_. See Supplementary File 2 for the full analysis at all time points.

**Figure 5.**
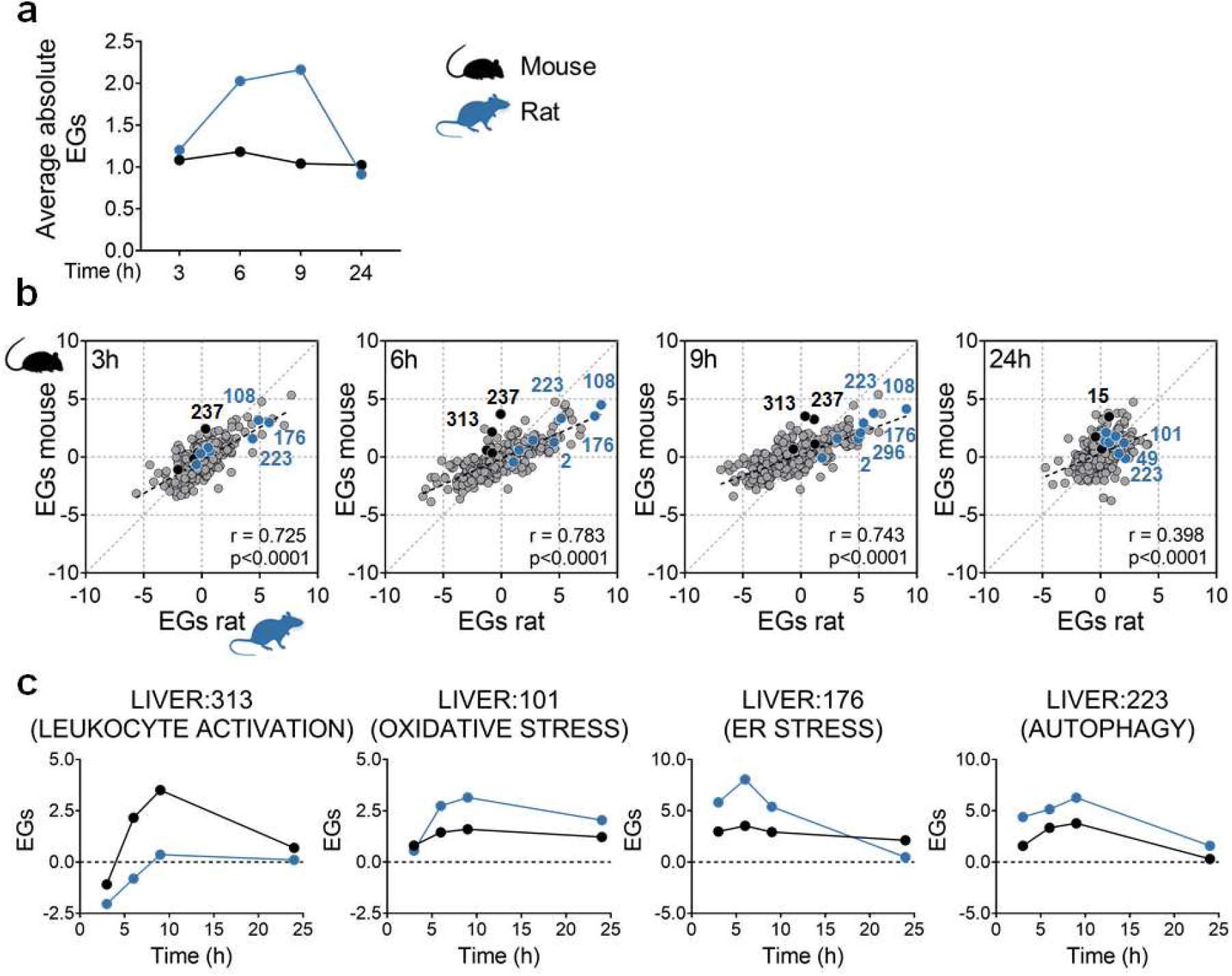
Rats show a more robust overall transcriptional response to equivalent NAPQI insult than mice. (**a**) Average absolute eigengene score (EGs) in APAP-treated mice and rats, based on WGCNA interrogation of the liver transcriptomics data. (**b**) Correlation plots of module EGs in the two species at the indicated time points. Pearson correlation coefficient (r) and P-value for each comparison are shown in the plot. EGs for relevant modules that are higher in mouse are depicted in black (LIVER:237, negative regulation of cell proliferation; LIVER:313, leukocyte activation; LIVER:15, cell death), whereas those higher in rat are depicted in blue (LIVER:108, LIVER:101, and LIVER:49, oxidative stress response; LIVER:176, endoplasmic reticulum stress response; LIVER:223, LIVER:2, and LIVER:296, autophagy). (**c**) EGs comparison across species for selected modules. See Supplementary Fig. 10 and Supplementary File 3 for more details.

To confirm that the apparent species differences in response to the equivalent NAPQI insult were preserved at the protein level, quantitative proteomics analysis was carried out on liver tissues collected from the vehicle and APAP-treated animals. Given the time required for transcriptional responses to manifest in altered protein expression levels, we focused on the samples collected 6, 9 and 24 h after dosing, and performed Ingenuity Pathway Analysis (IPA) to understand the pathway level effects of the observed protein expression changes. Liver tissue from APAP-treated mice showed activation of toxicity functions associated with liver injury and cell apoptosis, whereas liver tissue from rats exhibited stimulation of hepatocyte proliferation processes, further supporting the different extents of adaptation between the two species (Fig. 6a). Consistent with the observed gene level changes, liver tissue from APAP-treated rats showed a lower increase in the expression of proteins involved in inflammatory and immune responses (Fig. 6b), and a greater increase in the expression of proteins involved in the Nrf2 oxidative stress response and autophagy (Fig. 6c-d, and Supplementary Fig. 11), compared to mice. The altered expression of representative proteins from these processes (NQO1, HMOX1, SQSTM1 and LC3B) was confirmed at each time point in the two species via immunoblotting (Fig. 6e-j). Notably, in the case of LC3B, which is commonly used as a marker of autophagy ^22^, both the cytosolic LC3B-I and autophagosome-bound LC3B-II were expressed at higher levels in the livers of rats at the 0 h time point (i.e. following the 16 h fasting period) (Fig. 6i-j). In addition, whilst APAP induced autophagy at early time points in both species, as evidenced by both a decrease in abundance of SQSTM1 (at 3 h) and the conversion of LC3B-I to LC3B-II (at 3 and 6 h), the level of LC3B-I was found to be consistently higher in the rat across the timeframe of the study (Fig. 6h-j), suggestive of a greater autophagic capacity. Together, these data support a more robust activation of adaptive stress response pathways in rats compared with mice following exposure to an equivalent NAPQI insult.

**Figure 6.**
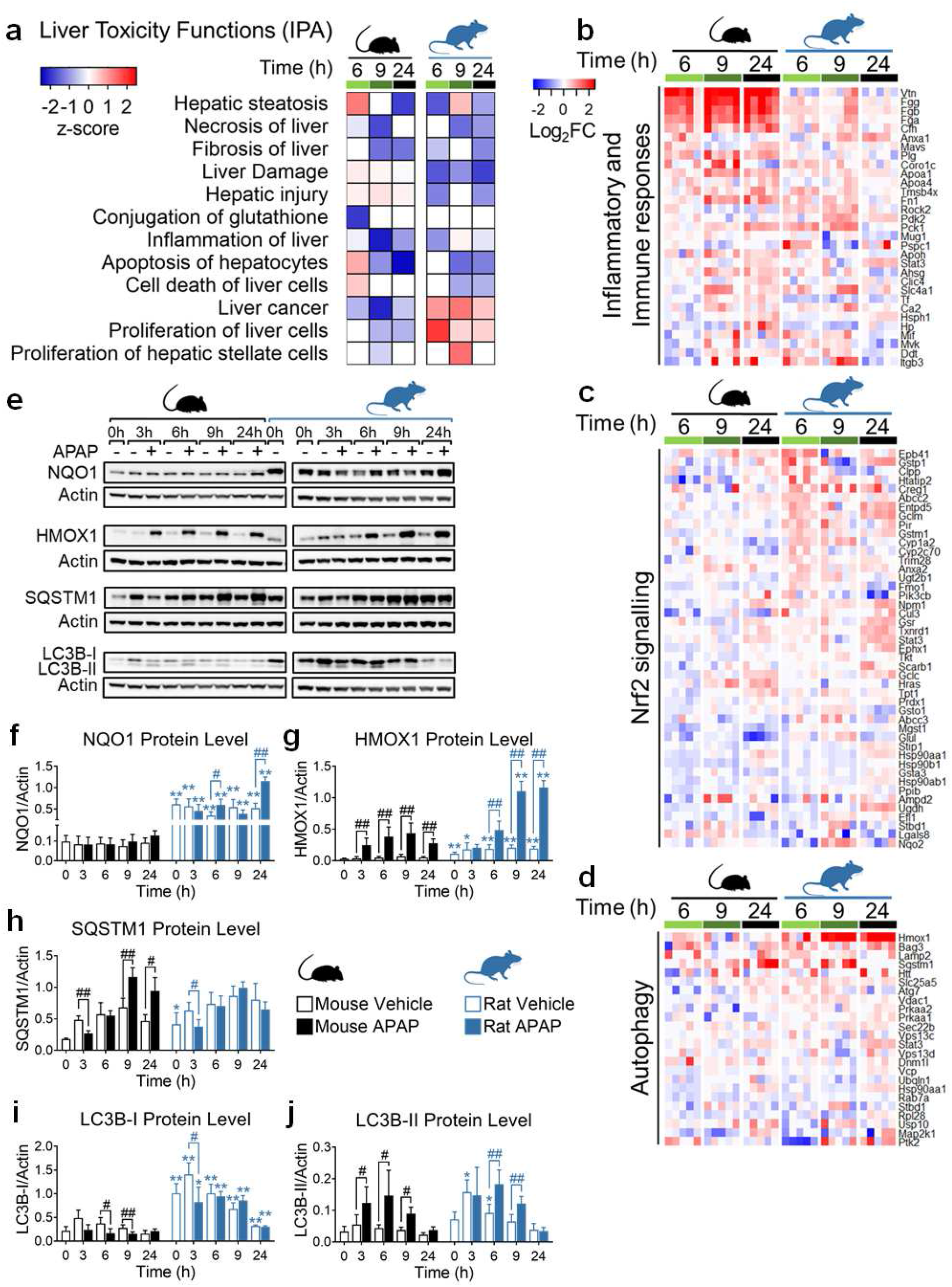
Proteomic analysis corroborates the more robust activation of adaptive stress response pathways in the liver of APAP-treated rats, compared with mice. (**a**) Comparative toxicity analysis (IPA-Tox) in the liver of mice and rats treated with 300 mg/kg and 1000 mg/kg APAP, respectively (n=5) at 6, 9, and 24 hours (SWATH proteomics). For each function a z-score was calculated at each time point against time-matched vehicle control animals. Heatmap plots showing changes in the expression of proteins involved in (**b**) inflammatory and immune responses, (**c**) Nrf2-signalling, and (**d**) autophagy. In (**b-d**), data are expressed as log_2_ fold-change *versus* time-matched vehicle control animals. (**e**) Protein expression levels of NAD(P)H dehydrogenase quinone 1 (NQO1), heme oxygenase 1(HMOX1), sequestosome 1 (SQSTM1), and light chain 3 isoforms B-I (LC3B-I) and -II (LC3B-II) at the indicated times after APAP or vehicle. Samples are pooled from 5 animals per time point. (**f-j**) Densitometric analysis of immunoblots. Protein levels were normalised to β-actin. Values are mean ± SD of all individual samples (n=5). Mann-Whitney U test. P-values are denoted as *p<0.05, and **p<0.01, comparison of mouse versus rat for each condition/time point, or ^#^p<0.05, and ^##^p<0.01, comparison of APAP-treated animals versus vehicle controls, as indicated.

### Rats exhibit a higher basal hepatic expression of stress response pathway markers compared with mice and humans

Whilst our ‘omics analyses highlighted a clear difference between mice and rats in the inducible activity of several stress response pathways, the timeframe for altered expression of the associated proteins (> 6 h) relative to the rapid onset of liver injury (> 2 h) means that these adaptations could not have occurred swiftly enough to directly influence the different sensitivities of the species to the toxic effects of the APAP OEDs. Given the limitations of probe-based microarray and qPCR data sets for comparing absolute gene expression levels across species, we used publicly available RNA-Seq (reads per kilobase million) data ^23^ to assess the basal expression levels of relevant genes in liver tissue from mice and rats, from birth to adulthood. This analysis highlighted the generally higher basal expression levels of a number of genes associated with the Nrf2 antioxidant and endoplasmic reticulum stress responses, as well as autophagic capacity, in rats compared with mice (Fig. 7). These findings were supported by our proteomics data; grouping the normalised expression quantities of orthologous proteins from the 0 h control animals into ranked bins confirmed the trend of higher basal expression of Nrf2 and autophagy-associated proteins in the livers of rats, compared with mice (Fig. 8, Supplementary File 4). Importantly, using the public RNA-Seq data, we also found that many genes associated with these stress response pathways were expressed at a higher basal level in the liver of rats than in humans, which overall better resembled mice (Fig. 7). We confirmed this at the protein level by immunoblot analysis of 0 h control tissues from animals in the APAP study, along with background liver tissue donated by eight patients undergoing elective resection of hepatic tumours (Fig. 9a-b). Notably, the latter samples exhibited marked inter-individual variability in the expression levels of the analysed proteins (Fig. 9a-b). Taken together, these data indicate that rats exhibit a greater basal and adaptive capacity of hepatic stress responses than mice and humans.

**Figure 7.**
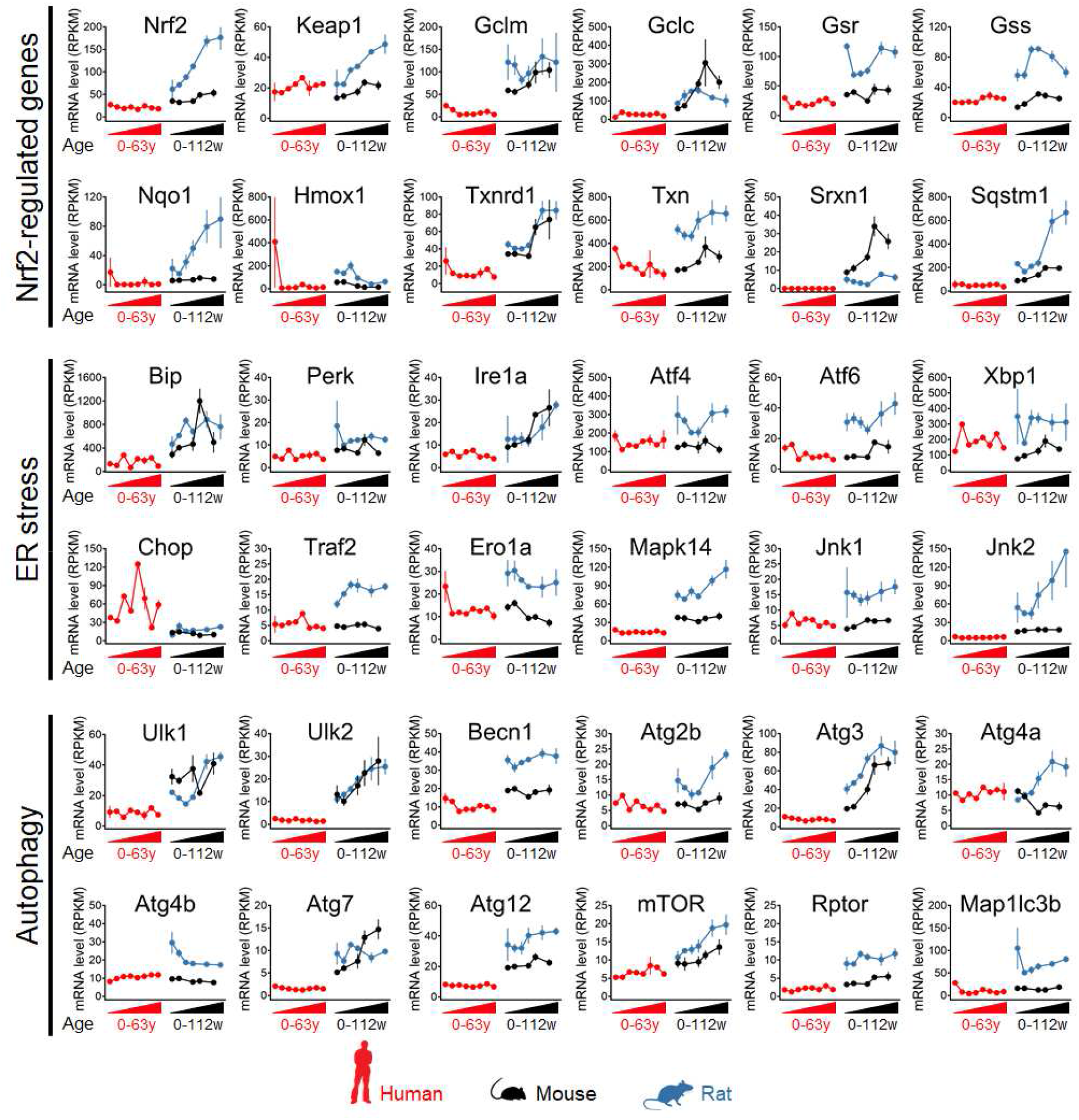
Higher basal expression levels of stress response pathway genes in the liver of rats, compared with mice and humans. Differences between species in the basal expression of Nrf2 genes and genes modulating endoplasmic reticulum (ER) stress and autophagy from birth to adulthood. mRNA levels are express as reads per kilo base per million (RPKM) ± SD ^23^. Human samples were grouped as follows: “neonates”, “infants” (6-9 months), “toddlers” (2-4 years), “school” (7-9 years), “teenagers” (13-19 years), “adults” (25-32 years), “middle-aged” (45-54 years), and “seniors” (58-63 years). Mouse samples (CD-1) were collected at week 0, 3, 14, 28, and 63, whereas rat samples (Sprague-Dawley) were collected at week 0, 3, 7, 14, 42, and 112.

**Figure 8.**
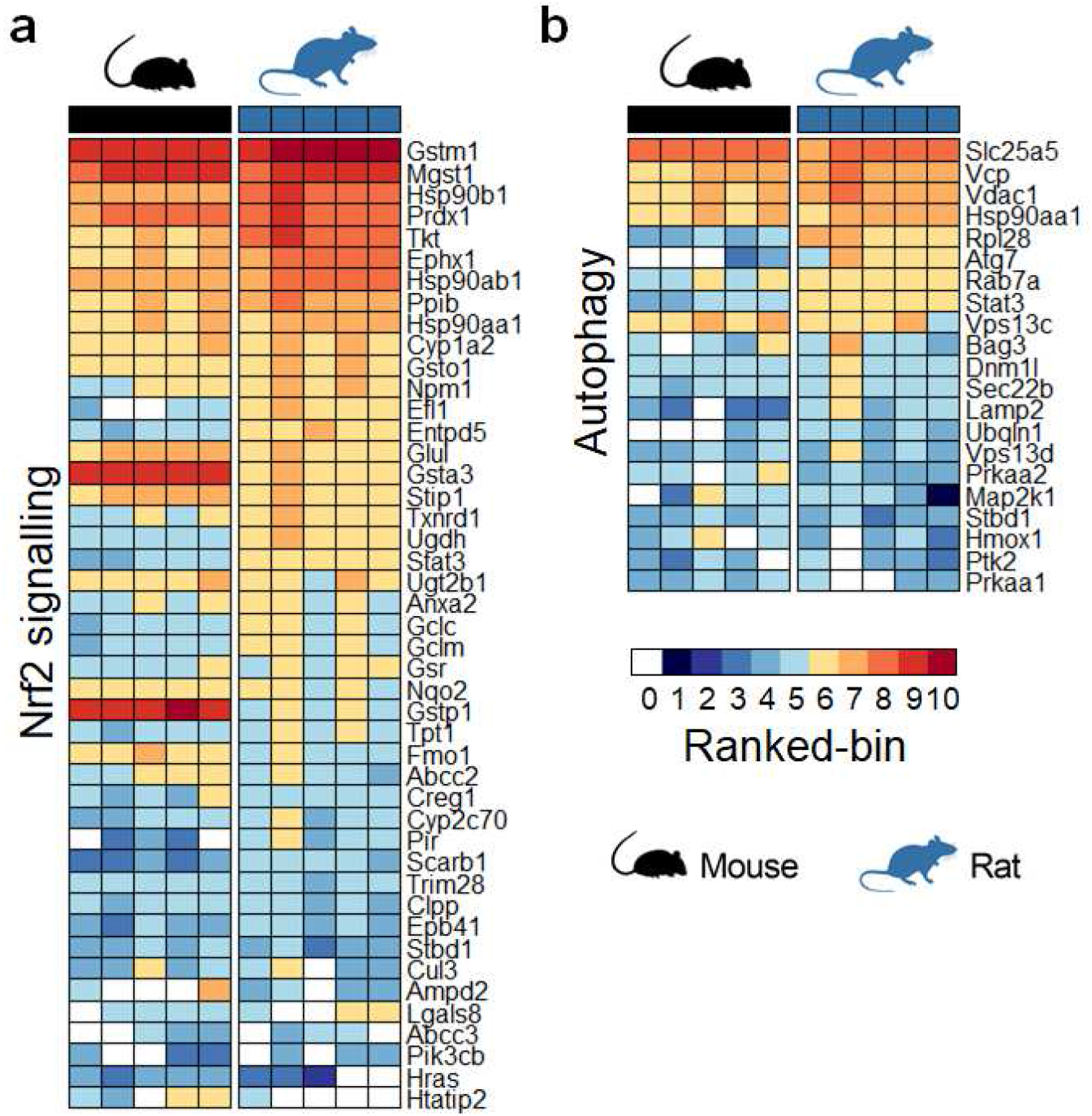
Higher basal expression levels of stress response pathway proteins in the liver of rats, compared with mice. Heatmaps showing basal expression levels of proteins associated with the (**a**) Nrf2 and (**b**) autophagy responses in the livers of untreated C57Bl/6J mice and Sprague-Dawley rats (0 h controls, n=5/species). Log_2_ transformed normalised protein expression values from SWATH proteomics were ranked and grouped into 10 bins, wherein proteins with the lowest abundance were assigned to bin 1, and those with the highest abundance to bin 10. A bin value of 0 (depicted in white) was assigned to proteins that were not detected. The difference amongst consecutive bins is 1.49 ± 0.16 (log_2_ expression). See Supplementary File 4 for more details.

**Figure 9.**
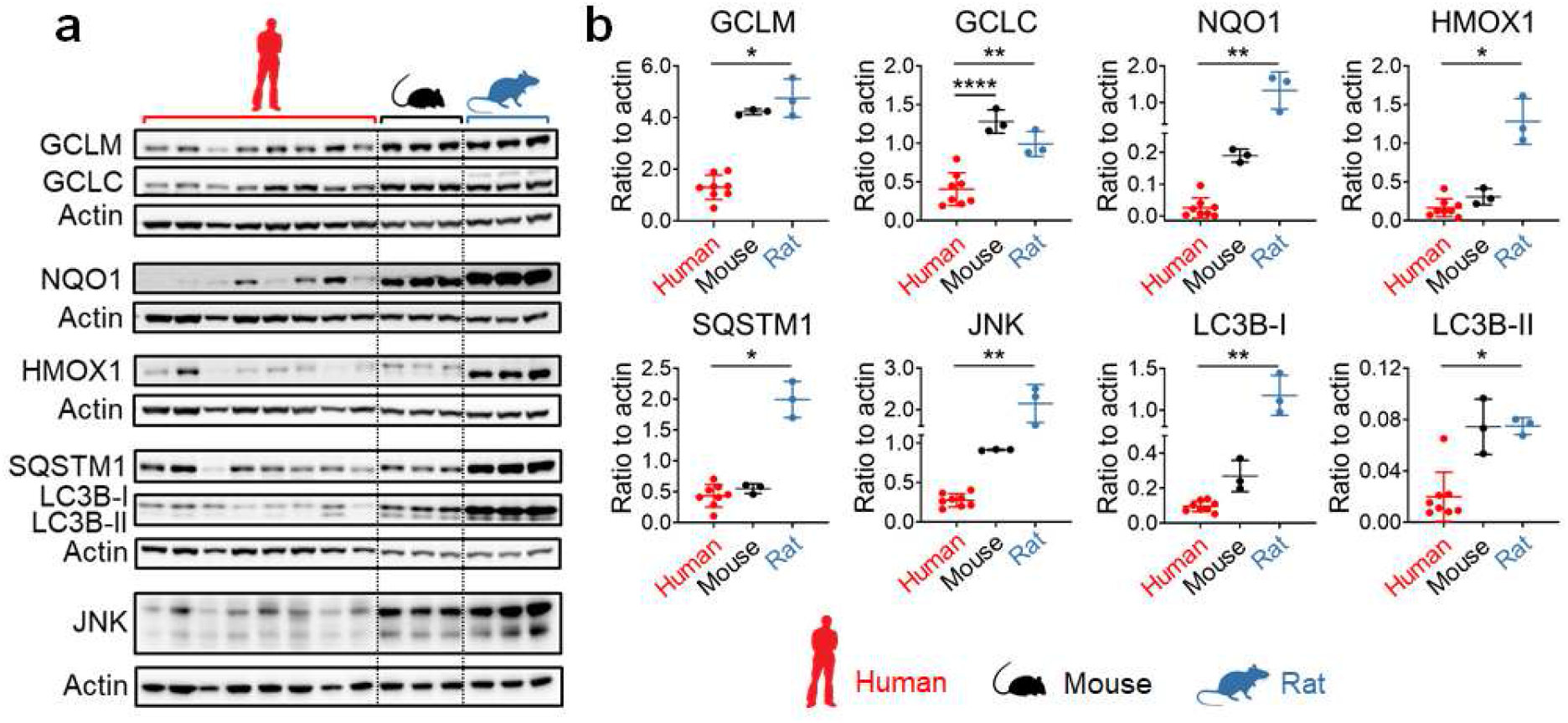
Confirmation of species differences in basal capacities of the Nrf2 and autophagy pathways. (**a**) Protein expression levels and (**b**) densitometric analysis of GCLM, GCLC, NQO1, HMOX1, SQSTM1, JNK, and LC3B isoforms I and II in liver tissue collected from patients undergoing liver resections (n=8) or untreated C57Bl/6J mice (n=3) and Sprague-Dawley rats (n=3). Protein levels were normalised to β-actin. Values are mean ± SD. One-way ANOVA or Kruskal-Wallis test, as appropriate. P-values are denoted as *p<0.05, **p<0.01, and ****p<0.0001, comparison as indicated.

## DISCUSSION

In the case of small molecule drugs, which make up a significant proportion of medicines submitted to global regulatory authorities each year, preclinical toxicity studies are expected to be performed in both a rodent and non-rodent species. Although the rat is the most commonly used rodent species, several studies have concluded that it may not be the most predictive of human drug toxicities ^1, 3, 24^, highlighting the need to better understand translation of safety data across species. Differences in metabolism and disposition are known to contribute to differences in species sensitivity to drug toxicity. However, less attention has been paid to the possibility that tissue properties, such as stress response capacity, may also contribute to cases of differential species susceptibility. We have used a combination of PBPK modelling and simulation coupled to multi-omics analysis to determine if differential sensitivity to APAP-induced liver injury in mice and rats is related to differences in the tissue response to equivalent burdens of the reactive intermediate NAPQI. Our study highlights the higher basal and inducible activity of the Nrf2 oxidative stress and autophagy responses in the livers of rats, compared with mice and humans. Our findings have important implications for species selection in preclinical safety testing of new medicines. Indeed, the higher stress response capacity of the rat may partly explain why it often under predicts the sensitivity of other species, including humans, to certain drug toxicities, including those associated with reactive metabolite formation and other forms of chemical stress.

Rats are well known to be relatively resistant to APAP liver toxicity, compared with mice and other sensitive species ^7^, due in part to differences in the extent of drug bioactivation and detoxication. Indeed, Gregus et al. ^12^ previously demonstrated that mice excrete over 50 % of a dose of APAP as toxic bioactivation products (APAP-GSH and associated hydrolysis products) whereas rats excrete only 9 % of APAP via these routes. As a result of this and the greater excretion of detoxication pathway metabolites (APAP-glucuronide and -sulfate) in rats, the ratio of toxic bioactivation products versus detoxication pathway metabolites was found to be 10-fold higher in mice than rats ^12^. Consistent with these findings, Boobis et al. ^14^ reported that primary hepatocytes from mice and rats exhibit large differences in the extent of covalent binding and degree of toxicity following exposure to APAP in vitro. Notably, Tee et al. ^13^ provided evidence that rat hepatocytes are relatively resistant to depletion of GSH and covalent protein binding upon direct exposure to NAPQI in vitro, compared with hepatocytes from hamsters, which are known to be highly sensitive to APAP liver toxicity. To overcome these differences in metabolic fate of the drug and enable a balanced assessment of the tissue response of mice and rats to a comparable NAPQI insult, we used PBPK modelling and simulation to support the use of 300 and 1000 mg/kg APAP as OEDs yielding an equivalent total hepatic burden of NAPQI in mice and rats, respectively. Despite this, rats remained more resistant to APAP hepatotoxicity compared with mice.

McGill et al. ^7^ also used doses of 300 and 1000 mg/kg (albeit via intraperitoneal, rather than oral, administration) in their investigation of the biochemical events underlying the differential sensitivity of mice and rats, respectively, to APAP liver toxicity. Consistent with our findings, the authors reported a substantial increase in serum ALT activity and marked centrilobular necrosis in mice, with minimal responses observed in rats ^7^. Total APAP-protein adducts reached similar levels in the two species, further confirming the equivalent degrees of chemical insult with these doses, and leading the authors to conclude that there are downstream factors responsible for the difference in sensitivity of mice and rats to APAP toxicity ^7^. In mice, JNK has been shown to undergo early phosphorylation and translocation to mitochondria partly as a result of the initial oxidative stress provoked by APAP ^25, 26^. Here, we have shown that p-JNK accumulates in liver samples from both mice and rats, peaking 3 h after the administration of APAP OEDs. In their study, McGill et al. ^7^ demonstrated that APAP-protein adducts and p-JNK translocation were consistently higher in mitochondrial fractions from mouse liver, and that rats therefore do not develop the critical levels of mitochondrial dysfunction or oxidative stress associated with APAP toxicity in mice. In light of our findings, it is possible that this is due to the greater capacity of the rat liver to cope with the initial chemical insult and mitigate against the progression of downstream toxic events.

Consistent with our findings, Xu et al. ^27^ recently reported that mice better resemble humans in terms of the basal hepatic expression of oxidative stress-related genes, whilst rats better resemble humans in terms of the expression of genes associated with lipid metabolism. Transgenic Nrf2 null mice have been reported to exhibit greater susceptibility to a large number of liver toxins, including APAP ^28^, whilst Kang et al. ^29^ recently reported that the response of an Nrf2-associated gene set was typically higher in primary rat hepatocytes, compared with human, exposed to a large panel of liver toxic and safe drugs. Autophagy has been implicated in the pathogenesis of DILI ^30^ and shown to act as a defence mechanism in APAP hepatotoxicity by removing APAP adducts ^31^ and damaged mitochondria ^32^. Notably, APAP protein adducts have been shown to be cleared at a faster rate in the livers of rats ^33^ compared with mice ^34^. The Nrf2 pathway and autophagy are linked by the p62/SQSTM1 (sequestosome 1) protein, which acts as a cargo receptor for autophagic degradation of ubiquitinated targets and activates Nrf2 by interfering with its ability to bind to its inhibitor Keap1 ^35^. SQSTM1 expression is also transcriptionally regulated by Nrf2, suggesting that SQSTM1 forms part of a regulatory feedback loop in the Nrf2 pathway ^36^. Hence, the differential activity of these and other cytoprotective pathways is expected to influence species sensitivity to a range of chemical insults.

In keeping with our finding of differences in the basal expression of stress response pathway components across liver tissues donated by eight patients, Kang et al. ^29^ recently reported quantitative differences in the drug-induced response of Nrf2-associated genes amongst primary hepatocytes isolated from different human donors. In addition, Liu et al. ^37^ have provided evidence that the hepatic expression of Nrf2-regulated genes is altered in patients with certain liver diseases. Hence, whilst the stress response capacity of most humans likely sits within an average range, it is possible that some individuals have relatively higher or lower abilities to respond to chemical insults, which could influence inter-individual sensitivity to some drug toxicities and the predictivity of relatively homogeneous animal models. Further work is required to fully understand the extent and consequences of such differences in stress response capacity between individuals.

A limitation of our in vivo study is that we have focused on only APAP as an exemplar hepatotoxic compound, and the C57Bl6/J and Sprague-Dawley strains of mouse and rat, respectively. Harrill et al. ^38^ have reported marked differences in sensitivity to the toxic effects of 300 mg/kg APAP across a panel of 36 inbred mouse strains, although C57Bl6/J mice were found to sit in the middle of this range, and the mechanism of APAP toxicity has been extensively investigated in this strain ^7^. In addition, our comparison of the basal expression of stress response pathway components used public transcriptomics data from CD-1 mice, alongside Sprague-Dawley rats ^23^. Furthermore, Monroe et al. ^39^ recently demonstrated that a large number of drugs that form chemically reactive metabolites, and were terminated due to clinical liver toxicity, provoke an activation of hepatic Nrf2 signalling, despite a lack of overt liver injury, in the rat (mixture of Sprague-Dawley and Han Wistar strains). Hence, whilst there will likely be some differences between strains, these observations indicate that, in general, the rat liver is highly resistant to chemical insult due to an increased capacity to respond to a range of stresses. Another limitation is that our study has focussed on the liver, which is one of the most common targets of drug toxicity both preclinically and in patients ^40^. However, a broader understanding of the degree of conservation of stress responses and other key physiological traits in different organ systems across preclinical models and humans is vital to inform the selection of appropriate species for preclinical testing to best reflect the clinical setting. Such knowledge will also be useful for the parameterisation of computational tools that are being designed to support risk assessment as a new drug candidate transitions from the preclinical to the clinical sphere ^41^, therefore improving the efficiency of the drug development process and maximising patient safety.

## METHODS

### PBPK modelling

Physiologically-based pharmacokinetic (PBPK) models were constructed using the Simcyp Animal simulator (v18r2; Certara Inc.) to predict the disposition of APAP and its metabolites arising from sulfation, glucuronidation, and cytochrome P450 mediated metabolism and glutathione conjugation of the resultant reactive-metabolite, NAPQI. Details of the animal PBPK simulators and the quality assurance system used in the development of each version of the Simcyp simulator have been described previously ^42–45^. The depletion of the phosphoadenosine 5’-phosphosulfate (PAPS) co-factor is a critical mechanism in accurately capturing the kinetics of APAP and its metabolites at higher doses. At higher doses the availability of PAPS becomes rate limiting resulting in an increased fraction of the APAP dose being metabolised through cytochrome P450 mediated metabolism producing NAPQI. A simple PAPS cofactor turnover/depletion model was implemented using the Lua ^46^ custom scripting facility implemented in Simcyp. This PAPS turnover model returns a scaling factor applied to intrinsic hepatic clearance through the sulfation pathway and capturing the shift in fraction metabolized at higher APAP concentrations.

Models were parameterised and their predictive performance verified using the observed data from the in vivo study reported here; exemplar simulations are provided in Supplementary Fig. 12. The PBPK models were used to perform simulations to determine APAP OEDs (mg/kg body weight). Here the OED is defined as the orally-administered APAP dose resulting in a total (cumulative over 24 h) hepatic NAPQI burden equivalent between the species of interest. Predicted OEDs were used as doses in the in vivo time course study described below. Data from this study was then analysed to determine the total hepatic NAPQI burden in each species. Based on glutathione depletion, total hepatic NAPQI burden was calculated by determining the loss of GSH based on the average GSH level (nmol) in control animals at 24 h (μ_GSH,vehicle_24h_) and the minimum GSH level (min_GSH,APAP_) in APAP dosed animals; this was then converted to a total hepatic burden (mg/g liver) assuming a 1:1 stoichiometry between NAPQI and glutathione (equation x). Since both treated and vehicle control animals were fasted for 16 h, GSH levels were below baseline at the start of the dosing period. We therefore assumed that GSH levels had returned to their respective baseline in both species after 24 h on resumption of feeding.

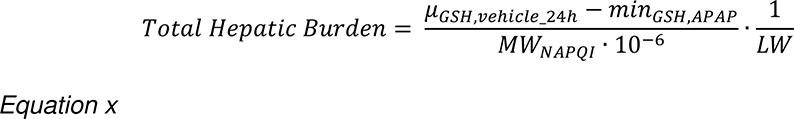

Liver weight (LW) was taken as 1.3 g and 9.0 g for mouse and rat, respectively, from Simcyp population library values (Simcyp v18r2; Certara Inc.); protein content was assumed to be 231.48 mg/g liver for both species ^47^. See Supplementary File 1 for further details of hepatic NAPQI burden calculations.

### Animals and dosing

Animal experiments were performed at Biologie Servier (France) in accordance with the European Council Directive 2010/63/EU on the protection of animals used for scientific purposes. The studies were approved by both Biologie Servier and TransQST consortium ethical review committees. For the dose-ranging study, male C57Bl/6J mice (8 weeks old; 15-22 g body weight; n=5 per dose) or Sprague-Dawley rats (5 weeks old; 119-144 g body weight; n=5 per dose) were fasted for 16 h prior to dosing. APAP (Sigma-Aldrich) was solubilised in 1 % (w/v) hydroxyethylcellulose (Sigma-Aldrich). Mice were administered 150, 300, 450 or 600 mg/kg APAP by oral gavage. Rats were administered 1000, 1300 or 1600 mg/kg APAP by the same route. Vehicle control groups were administered 1 % hydroxyethylcellulose. For the time course study, fasted mice (n=5 per time point) and rats (n=5 per time point) were administered 300 mg/kg and 1000 mg/kg APAP, respectively, or vehicle, by oral gavage. At relevant time points, the animals were sacrificed by exsanguination under isoflurane anaesthesia. Liver tissue and blood samples were collected at baseline (0 h) and at 3, 6, 9 and 24 h after administration of APAP or vehicle for the analyses described below. Given that the levels of serum markers of liver injury were significantly elevated in mice at the earliest 3 h time point, a separate in vivo study was conducted in mice administered 300 mg/kg APAP (by oral gavage) to collect samples at earlier time points (0.5, 1, 2, and 3 h, n=5-7) to better reflect the initial increase in serum markers of liver injury.

### Serum biomarker analysis

Blood samples were collected from a jugular vein or abdominal aorta of isoflurane-anaesthetised animals into tubes with a serum separator. Blood samples were centrifuged at top speed after clotting and analysed using a Cobas c501 analyzer (Roche). Alanine aminotransferase (ALT) and aspartate aminotransferase (AST) activities were measured kinetically at 37 °C according to the International Federation of Clinical Chemistry recommendations without Pyridoxal Phosphate (Roche). Total bilirubin concentration (TBIL) was measured using 3,5-dichlorophenyl diazonium (Roche).

### Liver glutathione content measurement

Liver tissue was homogenised in buffer (pH 7.4) comprising 143 mM NaH_2_PO_4_, 8 mM EDTA and 1.3 % sulfosalycilic acid, using an oscillating mill at 30 oscillations/sec for 3 min. The homogenates were then centrifuged (18,000 g for 5 min at 4 **°**C) and the supernatants collected. Total glutathione (GSH) was measured as described previously ^48^. The reaction was followed kinetically at 415 nm. Sample readings were interpreted against a GSH calibration curve and normalised to protein content, which was determined from the sample pellets (dissolved in 1 M NaOH at 60 **°**C) generated during the homogenisation step, using the bicinchoninic acid method (Sigma-Aldrich).

### Histopathological assessment

For each animal, the entire liver (gall bladder was carefully removed for the mouse) was rapidly excised and briefly rinsed in DNase/RNase -free distilled water. The left liver lobe was then separated and preserved in 10 % formalin. Fixed samples were mounted in paraffin wax, cut at approximately 4 µm in thickness, stained with haematin-eosin saffron (HES) and evaluated by two independent veterinary pathologists. The extent of centrilobular hepatocellular necrosis/degeneration was assigned a consensus semi-quantitative score as follows: 0, no degeneration/necrosis observed; 1, minimal degeneration/necrosis of individual or small groups of hepatocytes; 2, mild degeneration/necrosis that bridges some centrilobular zones; 3, moderate degeneration/necrosis that completely bridges centrilobular zones; 4, marked degeneration/necrosis that bridges centrilobular zones and extends into midzone areas; 5, severe degeneration/necrosis involving all hepatic lobular zones.

### Immunoblotting

Snap frozen liver tissue (30-40 mg) was homogenised in radioimmunoprecipitation buffer (Sigma Aldrich). Liver homogenate (20 μg) was separated by SDS-PAGE under reducing conditions, transferred to nitrocellulose membranes and subjected to immunoblot analysis as previously described ^35^. Details of all antibodies used in this study are provided in Supplementary Table 1. Immunoreactive bands were visualised using a ChemiDoc Imaging System (Bio-Rad) and volumes were quantified using Image Lab Version 6.1.0 (Bio-Rad), with normalisation to β-actin. Total protein normalisation was also used to exclude the effect of APAP treatment on β-actin expression, and yielded similar results (data not shown). For the representative blots, a pool of all samples for each treatment group and time point were loaded onto the gel. To allow for a fair comparison across species when mouse and rat samples were run on separate gels due to the number of time points/treatment conditions within an experiment, representative samples from the other species were added to each gel as internal controls, and the chemiluminescence signals associated with the two gels were quantified together.

### Transcriptomic analysis

Total RNA was isolated from liver tissue collected in RNAlater^®^ stabilizing reagent immediately prior to dosing (0 h) or at 3, 6, 9, and 24 h after administration of APAP or vehicle. Frozen liver samples were homogenised in Qiazol Lysis Reagent (Qiagen). Total RNA was purified following the manufacturer’s instructions. The RNA integrity in representative samples was verified using an Agilent BioAnalyzer. Total RNA (5 µg) was processed per the standard Affymetrix protocol for microarray target preparation. Resulting cRNA was fragmented and hybridized onto Affymetrix GeneChips, which were then washed, stained, and scanned using the standard Affymetrix procedure (Affymetrix GeneChip^®^ Mouse Genome 430 2.0 Array or Affymetrix GeneChip^®^ Rat Genome 230 2.0 Array). Quality control and low level analysis of raw fluorescence intensities was performed using the affy package (v1.64.0) in R v3.6.1 ^49^. Mouse data were processed using RMA normalization and BrainArray CDF Version 24 ^50^. Rat data were processed using RMA normalization and BrainArray CDF Version 19, which gives 100 % coverage on the DILI TXG-MAPr tool (https://txg-mapr.eu/) used for Weighted Gene Co-Expression Network Analysis (WGCNA) (manuscript in preparation). Differential expression analyses were carried out using the limma package (v3.42.2) in R ^51^. Orthologous genes between rat and mouse represented in the Rat Genome Database (RGD) were considered for comparison of the transcriptomic changes. Genes were considered differentially expressed when the adjusted P value (Benjamini-Hochberg correction) was less than 0.05 and there was at least a 1.5-fold change in expression. The full list of significant genes at each time point was analysed for membership in co-expressed gene sets using WGCNA, and an eigengene score (EGs, or module score) which summarises the log_2_ fold change of the respective constituent genes, was calculated for each module as previously described ^52^. The average absolute EGs was calculated by averaging the absolute scores across all the modules in each species at each time point. The list of the top 50 most significant genes in each module (ordered by significance) and their relative log_2_ fold-change can be found in Supplementary File 3.

### Pathway analysis

ClusterProfiler package (v3.18.0) ^53^ was used for gene set enrichment analysis (GSEA) on the lists of differentially expressed genes (DEGs) ranked by log_2_ fold-change. The number of permutations was set to 10,000. To remove differences in the setSize, genome wide annotation for Rat (package ‘org.Rn.eg.db’) was used for both species. Gene ontology (GO) biological processes ^54^ were considered significantly enriched with a FDR-corrected P_adj_< 0.05 and absolute Normalised Enrichment Score (NES) >1.5. The full list of significantly enriched GO biological processes in both species at all time points is provided in Supplementary File 2. Genes that were differentially expressed in either species at any time point within the lists of enriched genes in selected GO biological processes were used to generate heatmap plots showing changes at a gene level between species (data expressed as log_2_ fold-changes *versus* time-matched vehicle controls animals).

### qPCR analysis

RNA (500 ng) was reverse transcribed using a LunaScript RT SuperMix kit (New England Biolabs) according to the manufacturer’s instructions in a total 20 µL reaction mix. qPCR was performed using Luna Universal qPCR Master Mix (New England Biolabs) on an ABI ViiA-7 Thermocycler (Applied Biosystems). Primer sequences for mouse and rat genes are detailed in Supplementary Table 2. For relative quantification, data were analysed using the 2^-ΔΔCt^ method. The selection of GAPDH as the endogenous normalization control was based on analysis of the transcriptomic data using NormFinder ^55^, which ranked homologous genes with a coefficient of variation < 1% across treatments and time points, based on the calculated stability values.

### Proteomics sample preparation

Liver tissue (100-200 mg) was lysed by sonication on ice in 300 µL of 7 M urea, 2 M thiourea, 40 mM tris (pH 7.5) and 4 % w/v CHAPS buffer. After centrifugation (14,000 rpm for 10 min at 4 °C), the supernatant was collected and protein concentration was determined by Bradford assay. Next, 200 µg protein was reduced with 5 mM dithiothreitol at 37 °C for 30 min, alkylated with 0.15 M iodoacetamide, and incubated at 37 °C for 3 h with Trypsin/Lys-C (Promega) at a 25:1 protein:protease ratio (w/w). The reaction mix was then diluted 1:10 with 50 mM ammonium bicarbonate in order to reduce the urea concentration to less than 1 M, and incubated overnight at 37 °C. Peptides were then diluted to 5 mL with 10 mM potassium dihydrogen phosphate/25 % acetonitrile (ACN) and acidified to pH ≤ 3 with phosphoric acid prior to cation exchange chromatography. In order to generate the reference spectral libraries, a pool of 20 samples/species representative of all the experimental conditions was prepared. An aliquot of 1.5 mg of the pool was pre-fractionated on a Polysulfoethyl A column (200 × 4.6 mm, 5 μm, 300 A; Poly LC) at a flow rate of 1 mL/min. For individual samples, detergents and undigested protein were removed using a Bio-Scale Mini Macro-Prep High S 1 mL cartridge (Bio-Rad) on an Agilent 1100 HPLC system at 1 mL/min. The mobile phases consisted of 10 mM KH_2_PO_4_/25 % ACN w/v, pH 3 (phase A), and 10 mM KH_2_PO_4_/1 M KCl/25 % ACN w/v, pH 3 (phase B). A gradient from 0-50 % B in 90 min was applied to the Polysulfoethyl A column. Peptides were eluted from the cartridge using a gradient time program set as follows (phase B): 0 %; 0-0.5 min, 0-15 %; 0.5-5 min, 15 %; 5-5.10 min, 15-50 %; 5.10-7 min, 50 %; 7-7.10 min, 50-0 %; 7.10-10 min, 0 %. The system was then washed with 100 % phase A for 5 min after each sample to prevent sample carryover. Fractions of 1 mL were dried in a SpeedVac Concentrator Plus (Eppendorf), then reconstituted in 1 mL of 0.1 % trifluoroacetic acid and desalted using an Agilent Macroporous Reversed-Phase C18 column (4.6 × 50 mm mRP-C18, Agilent) on a 1260 Infinity HPLC system (Agilent).

### Data-dependent acquisition (DDA) and data analysis

Desalted fractions were reconstituted in 0.1 % formic acid and 0.5-1 μg of sample was loaded onto a UPLC Symmetry C18 nanoAcquity Trap Column (Waters). After a 10 min wash with 2 % ACN/0.1 % formic acid, the trap was switched in-line with a peptide BEH C18 nanoAcquity column (1.7 µm, 75 µm x 250 mm; Waters) and a gradient of 2-50 % ACN/0.1 % formic acid was applied over 120 min at a flow rate of 300 nL/min. DDA was performed on a Triple TOF 6600 (Sciex) in positive ion mode with a target of 25 MS/MS per cycle (2.8 sec cycle time), exclusion of former target ions for 20 sec, and using a mass range of 400-1800 for MS and 100-1500 for MS/MS. The data were searched using ProteinPilot 5.0 (Sciex) and the Paragon algorithm (Sciex) against the UniProtKB database (Mus musculus UP000000589, gene count 22,001, 55,366 proteins, last modified March 2021; Rattus norvegicus UP000002494, gene count 21,588, 29,934 proteins, last modified March 2021). Proteotypic peptides with no modifications except carbamidomethylation of cysteine residues were included in the library, so a ‘rapid’ search of the data was performed using ProteinPilot. Mass tolerance for precursor and fragment ions was 10 ppm. An FDR of 1 % was applied using the reversed database as decoy. This resulted in 5,253 and 5,823 proteins being included in the mouse and rat libraries, respectively.

### SWATH acquisition and data analysis

The same sample loading and chromatography conditions as described above were used for the individual sample acquisitions. Sequential Window Acquisition of all Theoretical Mass Spectra (SWATH) acquisitions were performed using 100 SWATH windows of variable effective isolation width to cover a mass range of 400-1600 m/z. The total cycle time was 3.1 sec. To account for batch effects, a pool of 20 samples representative of all the experimental conditions and a yeast digest quality control (Promega) were included with the samples in each batch. SWATH data were aligned with the spectral libraries using DIA-NN (v1.8) ^56^ with default settings in ‘robust LC (high accuracy)’ mode and annotated using the reference proteomes (UP000000589 and UP000002494) downloaded as FASTA files. Mass tolerances were determined automatically in DIA-NN for each run separately (Unrelated runs option). Match-between-runs was enabled to re-process the same dataset using a spectral library generated from the data-independent acquisition (DIA) data. Only proteins identified with proteotypic peptides and protein q-value below 0.01 were considered. Normalisation and differential expression analyses were carried out using the DEqMS package (v1.8.0) ^57^ in R. Protein quantities were log_2_ transformed and normalised using the equalMedianNormalization function, and Limma batch effect correction was applied. Ingenuity Pathway Analysis software (IPA; Qiagen) was used to investigate enriched canonical pathways and toxicity functions (IPA-Tox) in the two species. Comparison analyses were performed to evaluate changes in the z-score for each function over time and across species and reveal biological pathways underlying toxicity-specific phenotypes. Since the mouse and rat SWATH datasets were analysed separately, log_2_ transformed normalised protein expression values of orthologous proteins from the 0 h control animals (untreated) were ranked and grouped into 10 bins within each dataset for comparison of basal protein abundances across species^58^. Proteins with the lowest protein abundance values were assigned to bin 1, whereas those with the highest abundance values were assigned to bin 10. Proteins that were not detected (NA, not available) were assigned a bin value of 0. See Supplementary File 4 for more details.

### Analysis of public RNA-Seq data

To assess basal expression levels of relevant genes in liver tissue from humans, mice and rats, publicly available RNA-Seq data from Cadoso-Moreira *et al.* ^23^ were used (https://apps.kaessmannlab.org/evodevoapp/). Hepatic mRNA levels (reads per kilobase per million, RPKM) in humans (ArrayExpress E-MTAB-6814), mice (E-MTAB-6798), and rats (E-MTAB-6811) were compared. Human samples were grouped as follows: “neonates”, “infants” (6-9 months), “toddlers” (2-4 years), “school” (7-9 years), “teenagers” (13-19 years), “adults” (25-32 years), “middle-aged” (45-54 years), and “seniors” (58-63 years). Mouse samples (strain CD-1, RjOrl:SWISS) were collected at week 0, 3, 14, 28, and 63, whereas rat samples (strain Holtzman Sprague-Dawley) were collected at week 0, 3, 7, 14, 42, and 112.

### Human liver tissue

Human liver tissue was obtained by qualified medical staff at Aintree University Hospital (Liverpool, UK). All patients donated tissue as part of planned liver resections (see Supplementary Table 3 for details). Written, informed consent was obtained from each patient. The study protocol was approved by the National Health Service North West–Liverpool Central Research Ethics Committee (11/NW/0327) and conformed to the ethical guidelines of the 1975 Declaration of Helsinki.

### Statistical analysis

Statistical analysis was performed using StatsDirect 3 Software. Normal distribution was assessed by the Shapiro-Wilk test. A two-tailed student’s unpaired t-test or one-way ANOVA was used if normality was indicated. For non-normally distributed data, a two-tailed Mann Whitney U or Kruskal Wallis test was applied. Results were considered significant if P≤0.05.

### Data availability

The transcriptomic data discussed in this publication have been deposited in NCBI’s Gene Expression Omnibus ^59^ and will be accessible through GEO Series accession number GSE205203 (https://www.ncbi.nlm.nih.gov/geo/query/acc.cgi?acc=GSE205203) upon publication of the manuscript.

The SWATH proteomics data have been deposited to the ProteomeXchange Consortium (http://proteomecentral.proteomexchange.org) via the PRIDE partner repository ^60^ with the dataset identifier PXD034438. Data will be publicly available upon publication of the manuscript.

## Supporting information

Supplementary Experimental Procedures Tables and Figures

Supplementary File 1 - Calculation of hepatic NAPQI burden

Supplementary File 2 - GSEA

Supplementary File 3 - WGCNA

Supplementary File 3 - Binned Protein Expression

## Acknowledgements

This work is dedicated to the memory of Professor B. Kevin Park, the founding coordinator of the TransQST consortium, who died during the preparation of this manuscript. We acknowledge the contributions of Biologie Servier staff to the in-life phases of this work (Sylvie Roussel, Anne Marchau, Christine Bubois, Nathalie Delory, Claudette Chevalier, Laura Bochet, Eric Jacquemin, Catherine Cador, Laurent Gindrey, Jean-Pierre Creusillet, Agathe Buvat), and of Technologie Servier staff to toxicokinetics and metabolic analysis (Nathalie Landerouin, Jacky Pothier,Laurence Launay, Laurence Proust). We also acknowledge the contributions of AbbVie staff to the RNA extraction (Clarissa Woody) and the microarray analysis (Rita Ciurlionis). We acknowledge use of the CDSS Bioanalytical Facility provided by Liverpool Shared Research Facilities, Faculty of Health and Life Sciences, University of Liverpool. Lastly, we acknowledge financial support from the TransQST consortium, which received funding from the Innovative Medicines Initiative 2 Joint Undertaking under grant agreement no. 116030. This Joint Undertaking receives support from the European Union’s Horizon 2020 research and innovation program and EFPIA. This work was also supported by the Medical Research Council (MRC) as part of the Centre for Drug Safety Science (MR/L006758/1).

## Author contributions

GR: study concept, experiments, data interpretation, manuscript drafting. RSY: experiments, data interpretation, manuscript draft. LAL: experiments. HC: experiments. REJ: experiments, data interpretation. SJK: experiments, data interpretation. CPF: experiments, data interpretation, manuscript draft. DR: experiments, data interpretation. IG: data interpretation. AHR: experiments. SWF: experiments. ARJ: data interpretation. GVDC: study concept. GS: experiments. HB: experiments, data interpretation. RJW: study concept. RLJ: experiments, data interpretation, manuscript draft. MJL: experiments, data interpretation. DC: data interpretation, manuscript draft. JLS: funding acquisition, study concept, data interpretation, manuscript draft. CEG: funding acquisition, manuscript draft. IMC: funding acquisition, study concept, data interpretation, manuscript draft. All authors critically reviewed and approved the final manuscript.

## Competing interests

The authors declare no competing interests.

### Abbreviations

ACN: acetonitrile
ALT: alanine aminotransferase
APAP: acetaminophen
AST: aspartate aminotransferase
AUC: area under the curve
BCA: bicinchoninic acid
CHAPS: 3-[(3-cholamidopropyl)dimethylammonio]-1-propanesulfonate
DDA: data-dependent acquisition
DEGs: differentially expressed genes
DIA: data-independent acquisition
DILI: drug-induced liver injury
EDTA: ethylenediaminetetraacetic acid
EGs: Eigengene score
ER: endoplasmic reticulum
FDR: false discovery rate
GAPDH: glyceraldehyde 3-phosphate dehydrogenase
GCLC: glutamate-cysteine ligase catalytic subunit
GCLM: glutamate-cysteine ligase modifier subunit
GFR: glomerular filtration rate
GO: gene ontology
GSEA: gene set enrichment analysis
GSH: glutathione
HES: haematoxylin-eosin saffron
HMOX1: heme oxygenase 1
HPLC: high-performance liquid chromatography
IPA: Ingenuity Pathway Analysis software
JNK: c-Jun N-terminal kinase
Keap1: Kelch-like ECH-associated protein 1
LC3B: microtubule-associated proteins 1A/1B light chain 3B
LD50: lethal dose 50
LW: liver weight
MW: molecular weight
NAC: N-acetyl cysteine
NAPQI: N-acetyl-p-benzoquinoneimine
NES: normalised enrichment score
NQO1: NAD(P)H quinone dehydrogenase 1
Nrf2: nuclear factor-erythroid factor 2-related factor 2
OED: oral equivalent dose
PAPS: phosphoadenosine 5’-phosphosulfate
PBPK: physiologically-based pharmacokinetic
p-JNK: phosphorylated c-Jun N-terminal kinase
RGD: Rat Genome Database
RPKM: reads per kilobase per million
SDS: sodium dodecyl sulfate
SQSTM1: sequestosome 1
SWATH: sequential window acquisition of all theoretical mass spectra
TBIL: total bilirubin
WGCNA: weighted gene co-expression network analysis.

