## Supplementary Experimental Procedures Tables and Figures for "A systems approach reveals species differences in hepatic stress response capacity"

*Pharmacokinetic analysis*

Plasma samples (collected in heparin-coated tubes and stored at -80 °C until analysis) were thawed at room temperature and centrifuged (3000 g for 5 min at 4 °C). Each sample (10 µL) was then spiked with 10 µL internal standard (8 µg/mL for APAP d4 and APAP Gluc d3, 9 µg/mL for APAP Sul d3 and 0.6 µg/mL for APAP NAC d5, APAP Cys d5 and APAP GSH d3; all in MeOH) or MeOH alone as blank. Proteins were precipitated from each sample by adding 180 µL MeOH, mixed (1500 rpm for 5 min), stored at -20 °C for 20 min, centrifuged (3000 g for 5 min at 4 °C) and the supernatants collected. Each sample was further diluted with 600 µL water, mixed (800 rpm for 2 min), centrifuged (3000 g for 5 min at 4 °C) and the supernatant used for injection into the UPLC-MS/MS system. The samples were analysed using an UPLC-I Class system (Waters) coupled to an API 4000 QTrap mass spectrometer (Sciex). Separation was performed using an Acquity UPLC HSS T3, 1.8 µm, 100 x 2.1 mm with a 0.2 µm pre-filter (Phenomenex) and a column oven set to 40 °C. The injection volume was 2 µL and flow rate 0.6 mL/min. The mobile phases and gradient programme are summarised in Supplementary Table 4. Mass spectrometric analysis was performed in either positive or negative ion mode, as detailed in Supplementary Tables 5 and 6.

*Feed comparison study*

All experiments were performed in accordance with a license granted under the UK Animals (Scientific Procedures) Act 1986 and were approved by the University of

Liverpool Animal Ethics Committee. All animals received humane care according to the criteria outlined in the Guide for the Care and Use of Laboratory Animals. Male C57Bl/6J mice (7-8 weeks old) and male Sprague-Dawley rats (7-8 weeks old) were supplied by Charles River (UK). After the acclimatisation period (1 week), animals (n=4/group) were assigned to either RM1 (RM1P-E-FG, Special Diet Services, UK) or A04 (DS-SAFE-A04, Safe, France) diet for 7 days. All animals received food and water *ad libitum*. At the end of the study, the animals were culled via exposure to a rising concentration of $CO_2$, and liver tissue was excised and snap-frozen for later processing. RNA was extracted using a Monarch total RNA miniprep kit (New England Biolabs, UK) following manufacturer' instructions. Input RNA was reverse transcribed and qPCR was performed as described in the main manuscript.

### SUPPLEMENTARY TABLES

**Supplementary Table 1.** List of antibodies used for western blot experiments.

| Antigen | Host | Dilution factor | Company | Cat. n. |
|---|---|---|---|---|
| **Gclc** (Glutamate-Cysteine Ligase Catalytic Subunit) | Rabbit | 1:5000 | Abcam (UK) | ab41463 |
| **Gclm** (Glutamate-Cysteine Ligase Modifier Subunit) | Rabbit | 1:5000 | Abcam (UK) | ab126704 |
| **Hmox1** (Heme Oxygenase 1) | Rabbit | 1:5000 | Abcam (UK) | ab13243 |
| **LC3B** (Autophagy marker Light Chain 3 isoform B) | Rabbit | 1:750 | Cell Signaling Technology (US) | #2775 |
| **Nqo1** (NAD(P)H dehydrogenase quinone 1) | Goat | 1:2000 | Abcam (UK) | ab2346 |
| **p-SAPK/JNK** (Phospho-stress-activated protein kinases/Jun amino-terminal kinases) | Rabbit | 1:1000 | Cell Signaling Technology (US) | #4668 |
| **SAPK/JNK** (Stress-activated protein kinases/Jun amino-terminal kinases) | Rabbit | 1:1000 | Cell Signaling Technology (US) | #9252 |
| **Sqstm1** (Sequestosome 1) | Rabbit | 1:2500 | Sigma-Aldrich (UK) | P0067 |
| β-actin | Mouse | 1:10000 | Abcam (UK) | ab6276 |
| Rabbit IgG | Goat | 1:5000 | Sigma-Aldrich (UK) | A9169 |
| Mouse IgG | Rabbit | 1:5000 | Sigma-Aldrich (UK) | A9044 |
| Goat IgG | Rabbit | 1:5000 | Agilent/Dako (US) | P044901-2 |

**Supplementary Table 2.** Details of specific mouse and rat primers used in real-time qPCR experiments.

| Gene | Primer | Mouse sequence (5'→3') | Rat sequence (5'→3') |
|---|---|---|---|
| **Abcb1a** | Fwd | CGGAGTCAGACAGAACAAGAAGA | CAGCATCGCCGAGAACATTG |
| | Rev | AATGCTTCCAGGCATAAGCG | CGACAGCTGAGTCCCTTTGT |
| **Abcc3** | Fwd | AAGACTCGGGTGCTGGTAAC | TCCCACCTGAGTGCATTTGT |
| | Rev | TGCAAGGCTGCTTCATGGTC | TCAAACTGGGTACGGATGGT |
| **Atg101** | Fwd | CTCGGTGCGGGTCCAAATAG | AGTGGGTCTTGACCGAACTG |
| | Rev | CCCTTCTTCCCTAACGCGAA | CGCAAACCAAGTGAGGTGAG |
| **Atg12** | Fwd | TAAACTGGTGGCCTCGGAAC | TGGGGATGAGCCACAAATGAA |
| | Rev | ATCCCCATGCCTGGGATTTG | AGTCTCTTCCCACAGCATCAA |
| **Atg14** | Fwd | ACTGCTGGGCTCGTGTTTTA | CCAGAGCGGTGATTTCGTCT |
| | Rev | TTGCTGTAGGCGGTAGTTGG | CGTTTTCCTTCCATGGCCTTT |
| **Atg16l2** | Fwd | AGAAGCTATGCACTGGCAGG | CAAAAGGCGCTCTTCTTGGAG |
| | Rev | CATTAACAGCAGTGCAGTGGG | TGGCTAGCAGTTCAGCCTTC |
| **Bbc3** | Fwd | ATAGAGCCACATGCGAGCG | AACTAGGTGCCTACACCCGT |
| | Rev | GTGGGTTGCTATTGAGGCAC | TGGTGCAGAAAAAGTCCCCC |

| | | | |
|---|---|---|---|
| **Edem1** | Fwd | GCGCTTCAAAATAATGCCCG | GGGTGTGTGTGAGGACGATT |
| | Rev | CCGAAGACCAACCAGAGCAC | GCAAACTTCCCATTCGCAGG |
| **Gapdh** | Fwd | TGTCCGTCGTGGATCTGAC | TTCAACGGCACAGTCAAGGC |
| | Rev | CCTGCTTCACCACCTTCTTG | TCACCCCATTTGATGTTAGCG |
| **Gclc** | Fwd | ATGATAGAACACGGGAGGAGAG | AAAGCTTGGCTTAATCTACAGTTCA |
| | Rev | TGATCCTAAAGCGATTGTTCTTC | GGGAATAGTCTGCATCCTGCTT |
| **Gclm** | Fwd | AATCAGCCCCGATTTAGTCAG | TCAAGCTCACAACTCAGGGG |
| | Rev | CGATCCTACAATGAACAGTTTTGC | CGCCTCAGTGACGCTTTTTG |
| **Hmox1** | Fwd | GTCAAGCACAGGGTGACAGA | GGAAAGCAGTCATGGTCAGTCA |
| | Rev | ATCACCTGCAGCTCCTCAAA | CCCTTCCTGTGTCTTCCTTTGT |
| **Hsf1** | Fwd | CAACTGCCTTCATTGACTCCA | CCTCCTGTGTGTTTCCTCCTG |
| | Rev | GGCTCCGGTTGTGTCCATAG | GCCAGATTGGTCCCCTAAAG |
| **Il1r1** | Fwd | TGCCTCCCAGTAAAACAGTC | CCTCTGCCTCTTGACGATGG |
| | Rev | TCTCTTCCCAATCCAGTTCC | TGGTATGTGTAGGACGTGCG |
| **Keap1** | Fwd | CACAGCAGCGTGGAGAGA | ATGTGATGAACGGGGCAGTC |
| | Rev | CAACATTGGCGCGACTAGA | AAGAACTCCTCCTCCCCGAA |
| **Lgals3** | Fwd | GAGTACTAGAAGCGGCCGAG | CAGTGCCCTACGATATGCCC |
| | Rev | CCTGATTAGTGCTCCACCCG | TGGGCTTCACTGTGCCTATG |
| **Mafg** | Fwd | ATGACGACCCCCAATAAAGGA | GCCTTAAAGGTGAAGCGGGA |
| | Rev | CACCGACATGGTTACCAGC | AGGTGCTGGTTCAACTCTCG |
| **Nle1** | Fwd | CGTTCTACGTCCACGATGCT | TGCTGAGATTGTGTCCTCGC |
| | Rev | CAGGTACTTTCCTGTGGGGC | GAAATGACGGCTTCGCTGTG |
| **Nqo1** | Fwd | TTTAGGGTCGTCTTGGCAAC | GTTTGCCTGGCTTGCTTTCA |
| | Rev | GTCTTCTCTGAATGGGCCAG | ACAGCCGTGGCAGAACTATC |
| **Nrf2** | Fwd | CATGATGGACTTGGAGTTGC | CTTGCTCTTGGGAACAAGGAA |
| | Rev | CCTCCAAAGGATGTCAATCAA | CGACAGAAACCTCCATCTTCTG |
| **Prkaa1** | Fwd | TCTTCTCCTTAACTCCTCCCTC | TTCGGGAAAGTGAAGGTGGG |
| | Rev | TGTGTGTGGCATTCCATTCATC | TCTCTCTGCGGATTTTCCCG |
| **Sesn2** | Fwd | ACTGCGTCTTTGGCATCAGA | TCCCCCTAAGCCTGTTCTGT |
| | Rev | GTCTTCTCAGGGTAGCAGGC | CCACCAGGCATGGGAGAAAA |
| **Slc25a37** | Fwd | GGGCCTGAACGTGATGATGA | CCCCTTCCAATCTATCCACTTC |
| | Rev | ACTCCCAGCTACCCCATTAG | TCCTGAGATGATGTGAGACTG |
| **Smad7** | Fwd | TTTCCTCCTCCTTTCTCGTC | TTTTTCCCCCCACCCTTCCAAC |
| | Rev | CAAAACACACACACACAACC | AACACACCACCTTCTCGCAC |
| **Socs3** | Fwd | CACAGCCTTTCAGTGCAGAGTA | CCCCGCTTTGACTGTGTACT |
| | Rev | CGTAAGAGCAGGCGAGTGT | AAAGGAAGGTTCCGTCGGTG |
| **Srxn1** | Fwd | ACTATTCCTTTGGGGGCTGC | ACCTCCTGATACCCCACTCC |
| | Rev | GCTTGGCAGGAATGGTCTCT | GGAACCCCCTCATTCTTGGG |
| **Tgfbr2** | Fwd | CACGTTCCCAAGTCGGATGT | CCCCCGTTTGGTTCCAGAGT |
| | Rev | GTTTCAGTGGATGGATGGTCCT | GGTCTCTCAGCACGTTGTCT |
| **Trib3** | Fwd | CCTGCGTGATGACTGGATCA | GGGACTCCGAGATAGGCTCA |
| | Rev | CCGCTTTGCCAGAGTAGGAT | AGATGTGGCTCGCATCTTGT |
| **Txnrd1** | Fwd | TGTCAGGACAGCCAGTACTCTG | ACATGGAATTGGAATTTGGGTT |
| | Rev | CGGTCATGTAACCTAGGAGCTAA | CAAAGGAGTGGACTGTTGAGTTT |

**Supplementary Table 3.** Details of patients donating liver tissue as part of planned liver resection.

| Donor ID | Sex | Age | BMI | Indication | Underlying liver disease |
|---|---|---|---|---|---|
| S217 | 78 | F | 30.3 | CCA | Mild macrovesicular steatosis |
| S200 | 70 | M | 29.9 | HCC | None |
| S005 | 60 | F | 28.7 | CCA | Moderate microvesicular steatosis |
| S006 | 48 | M | 24.0 | CCA | Non-cirrhotic fibrosis |
| S285 | 71 | M | 21.8 | CCA | None |
| S205 | 73 | F | 28.9 | CRLM | Mild macrovesicular steatosis |
| S002 | 56 | F | 21.1 | CCA | None |
| S201 | 69 | M | 32.4 | HCC | Non-cirrhotic fibrosis/macrovesicular steatosis |

BMI, body mass index; HCC, hepatocellular carcinoma; CCA, cholangiocarcinoma; CRLM, colorectal cancer liver metastases.

**Supplementary Table 4.** LC-MS/MS conditions for pharmacokinetic analysis of APAP metabolites in plasma. Details of the liquid chromatography conditions used for analyte separation.

| Time (min) | 0 | 0.5 | 1.85 | 1.90 | 2.50 | 4.0 | 5.0 | 5.1 | 6 | 6.1 | 8.1 |
|---|---|---|---|---|---|---|---|---|---|---|---|
| % A ($H_2O$ + 0.1 % HCOOH) | 95 | 95 | 93 | 92 | 90 | 84 | 75 | 5 | 5 | 95 | 95 |
| % B (Methanol + 0.1 % HCOOH) | 5 | 5 | 7 | 8 | 10 | 16 | 25 | 95 | 95 | 5 | 5 |
| Curve | - | 6 | 6 | 6 | 6 | 6 | 6 | 6 | 6 | 6 | 6 |

**Supplementary Table 5.** Mass spectrometric parameters used in either negative or positive mode.

| | |
|---|---|
| Ionization mode: | Positive in period 2 and negative in periods 1 and 3 |
| Probe: | Horizontal: 5mm, Vertical: 2mm, Protrusion: 0.5mm |
| Curtain gas (CUR): | Nitrogen, 35psi |
| Collision gas (CAD): | Nitrogen, Medium (negative) and 6 (positive) |
| Ion Spray voltage (IS): | -4500 V (negative) and 5500 V (positive) |
| Temperature (TEM): | 500 °C |
| Ion source gas 1(GS1): | Air, 50 psi |
| Ion source gas 2 (GS2): | Air, 50psi |
| Entrance potential (EP): | 10 V |
| Interface heater (ihe): | On |
| Period 1: | 2.574 min |
| Period 2: | 1.205 min |
| Period 2: | 4.250 min |
| Total Run: | 8.030 min |

**Supplementary Table 6.** MRM parameters for the absolute quantification of each APAP metabolite.

| Compounds | MRM Transition | Mode | DP (V) | CE (eV) | CXP (V) | Dwell (msec) | Retention times | Period |
|---|---|---|---|---|---|---|---|---|
| APAP Gluc | 326.1 >149.9 | Negative | -75 | -38 | -11 | 50 | 1.64 min | 1 |
| APAP Gluc d3 | 329.1 > 153.0 | Negative | -75 | -40 | -7 | 50 | 1.62 min | 1 |
| APAP Sul | 230.0 >149.8 | Negative | -70 | -28 | -9 | 50 | 1.98 min | 1 |
| APAP Sul d3 | 232.9 > 152.8 | Negative | -60 | -28 | -11 | 50 | 1.96 min | 1 |
| APAP Cyst | 269.0 > 181.9 | Negative | -65 | -26 | -13 | 50 | 2.30 min | 1 |
| APAP Cyst d5 | 274.0 > 186.9 | Negative | -60 | -22 | -9 | 50 | 2.27 min | 1 |
| APAP | 152.1 > 110.1 | Positive | 61 | 23 | 18 | 100 | 2.71 min | 2 |
| APAP d4 | 156.1 > 114.1 | Positive | 61 | 23 | 18 | 100 | 2.68 min | 2 |
| APAP NAC | 311.0 > 181.9 | Negative | -60 | -24 | -9 | 100 | 4.84 min | 3 |
| APAP NAC d5 | 315.9 > 186.9 | Negative | -60 | -24 | -9 | 10 | 4.82 min | 3 |
| *APAP GSH | 457.1 > 328.1 | Positive | 71 | 23 | 10 | 50 | 3.57 min | N/A |
| *APAP GSH d3 | 460.1 > 331.1 | Positive | 71 | 23 | 10 | 50 | 3.5 min | N/A |

*Metabolites analysed separately from the main run under positive mode during the entire LC-MS/MS run.

### SUPPLEMENTARY FIGURES AND FIGURE LEGENDS

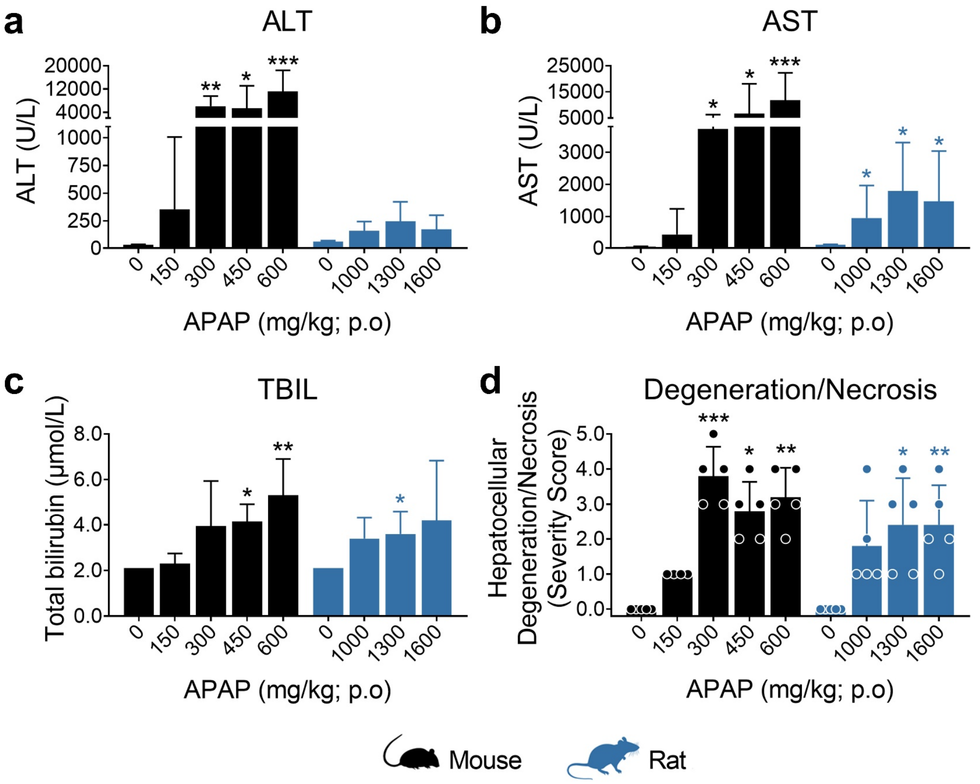


**Supplementary Figure 1.** Dose-ranging study to compare liver tissue responses of mice and rats to APAP. Male C57Bl/6J mice and Sprague-Dawley rats were administered 150-600 mg/kg or 1000-1600 mg/kg APAP, respectively, by oral gavage. We were unable to test higher doses in rat due to the limited solubility of the drug. Liver injury serum markers (ALT (**a**), AST (**b**) and TBIL (**c**)) and degeneration/necrosis (**d**) were evaluated 24 h after dosing. The extent of centrilobular hepatocellular degeneration/necrosis was assigned a semi-quantitative score as follows: 0 = none observed, 1 = minimal, 2 = mild, 3 = moderate, 4 = marked and 5 = severe (see main methods for full details). Values are mean ± SD (n=5). In (**d**) severity scores are shown for each animal. Statistical significance was determined between APAP-treated and vehicle control groups per species (Kruskal-Wallis with Dunn's multiple comparison test). P-values are denoted as *p<0.05, **p<0.01, or ***p<0.001.

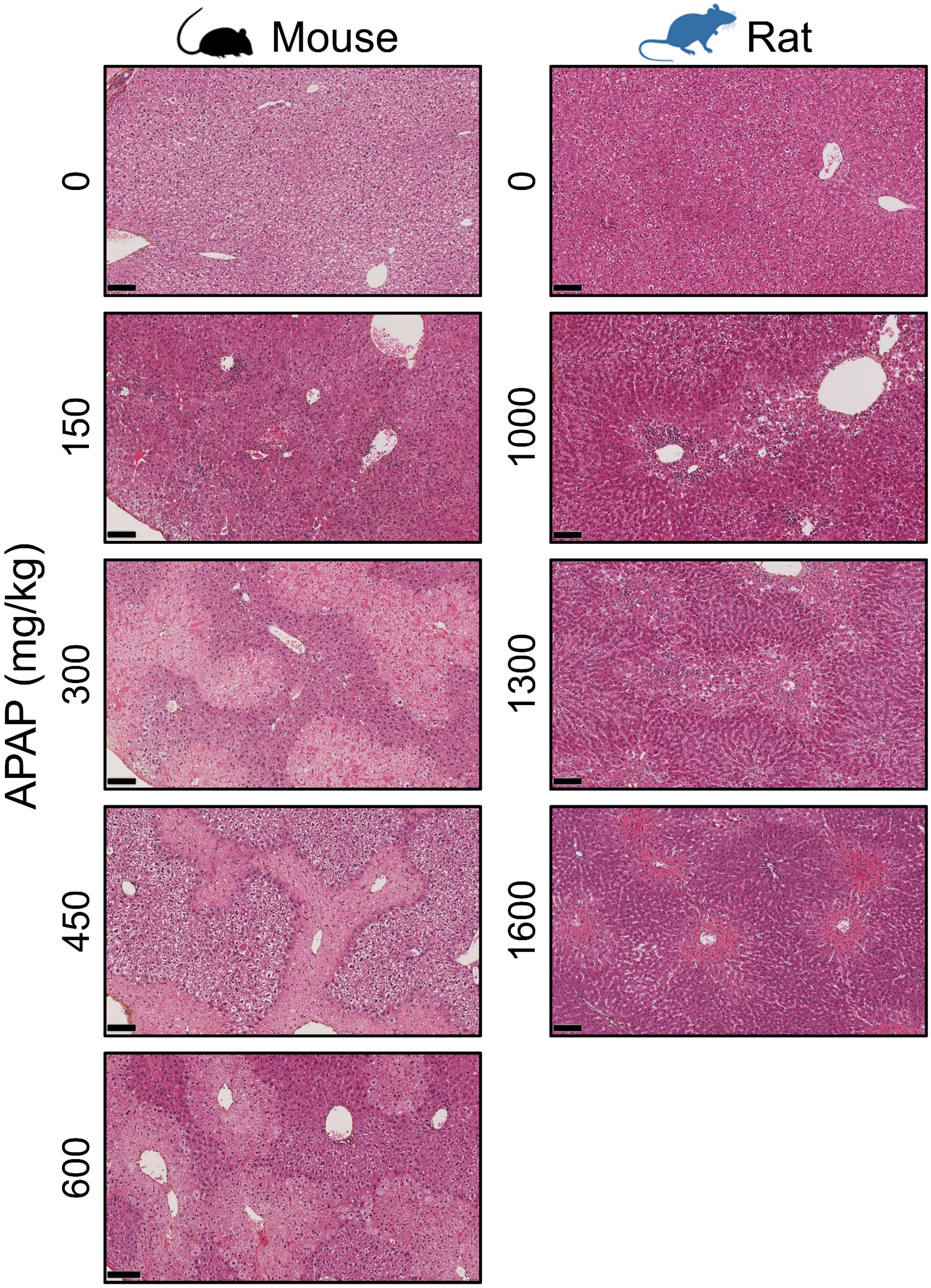


**Supplementary Figure 2.** Dose-ranging study to compare liver tissue responses of mice and rats to APAP. Male C57Bl/6J mice and Sprague-Dawley rats were administered 150-600 mg/kg or 1000-1600 mg/kg APAP, respectively, by oral gavage. Representative images of hematoxylin-eosin saffron stained liver sections from mice and rats treated with APAP at the indicated doses. Scale bar = 100 µm.

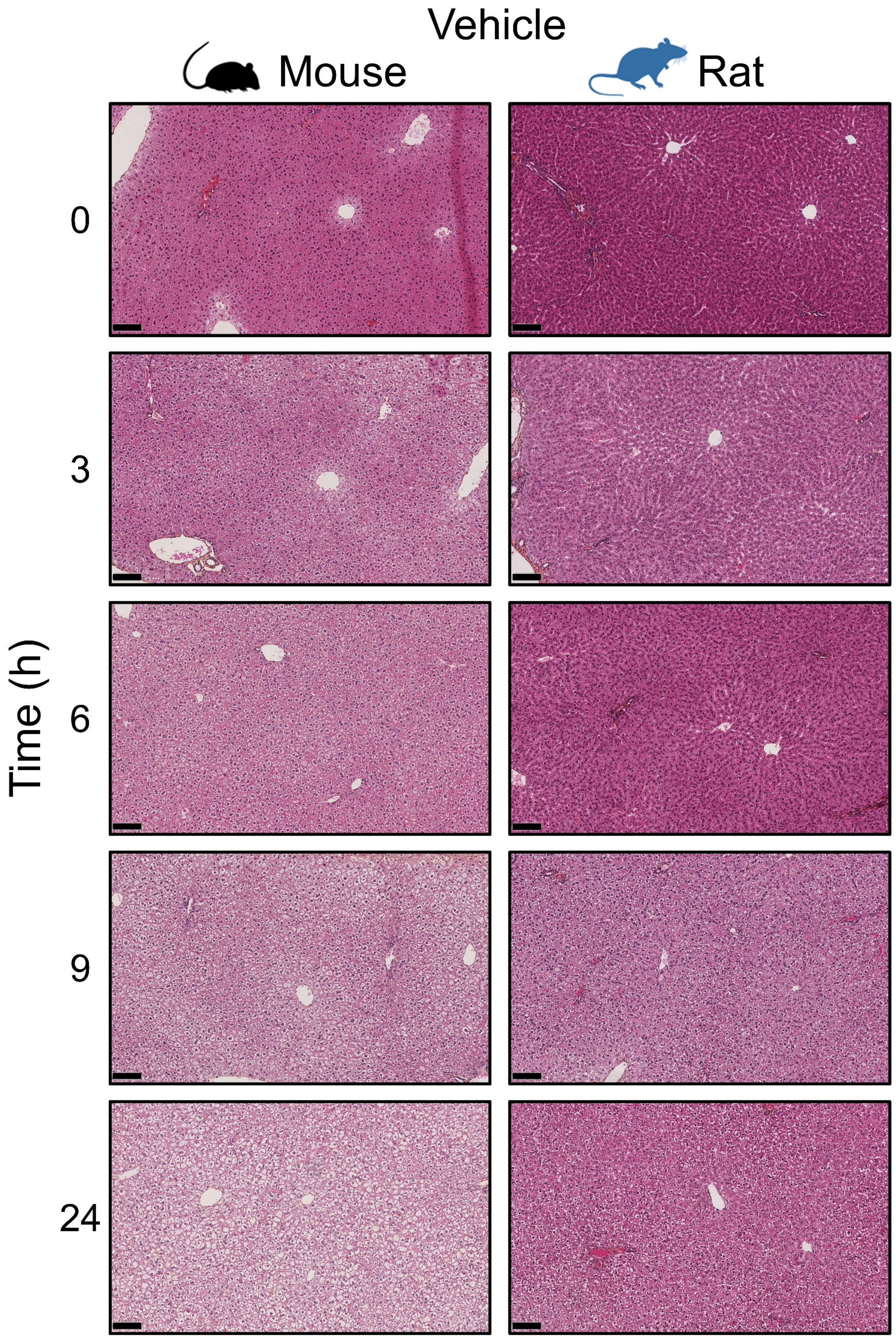


**Supplementary Figure 3.** Time course study to compare liver tissue responses of mice and rats to APAP. Representative images of haematin-eosin saffron stained liver sections from male C57Bl/6J mice and Sprague-Dawley rats at baseline (0 h) or treated with vehicle (1 % w/v hydroxyethylcellulose) by oral gavage at the indicated time points. Scale bar = 100 µm.

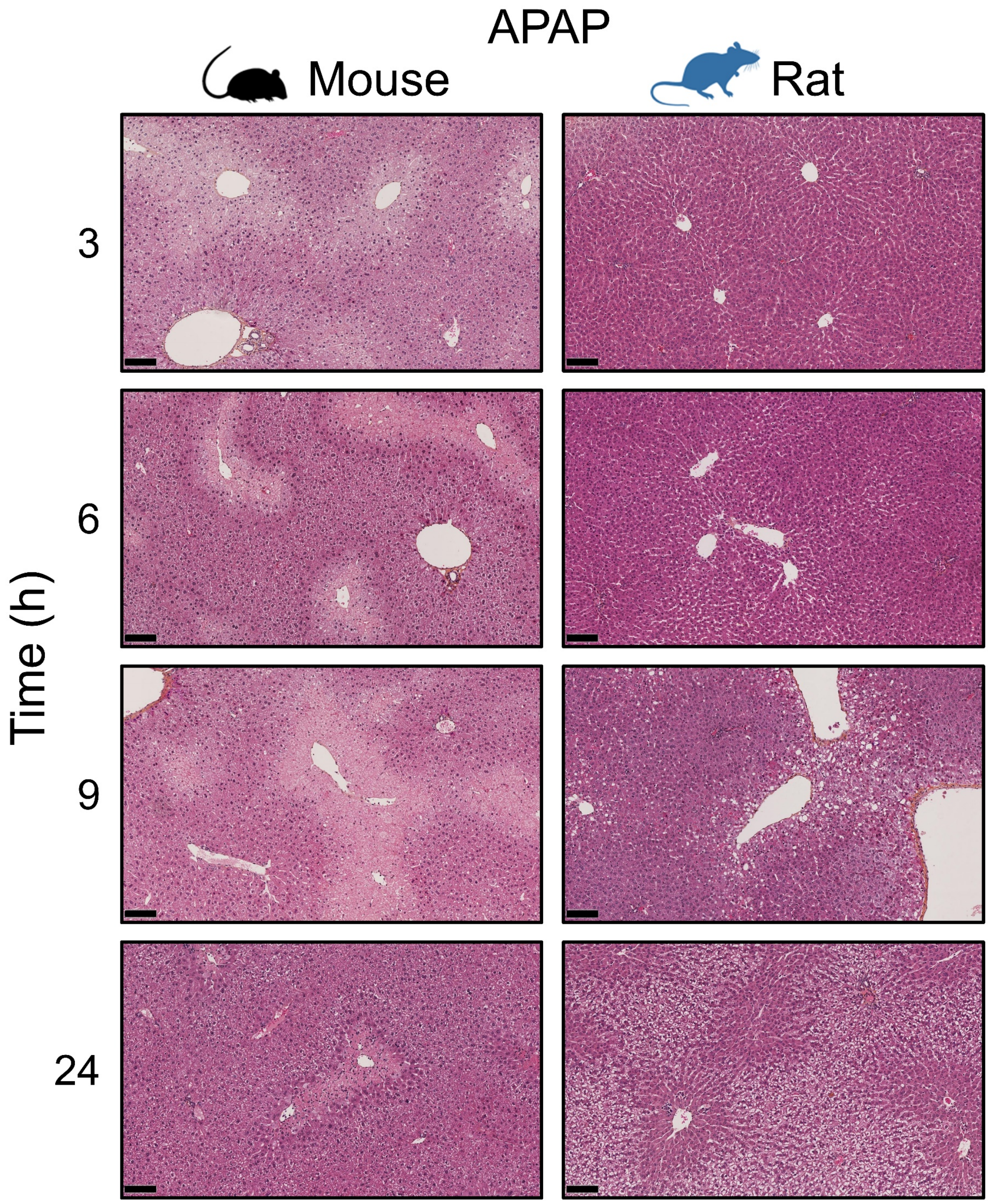


**Supplementary Figure 4.** Time course study to compare liver tissue responses of mice and rats to APAP. Representative images of haematin-eosin saffron stained liver sections from male C57Bl/6J mice and Sprague-Dawley rats treated with 300 mg/kg and 1000 mg/kg, respectively, by oral gavage at the indicated time points. Scale bar = 100 µm.

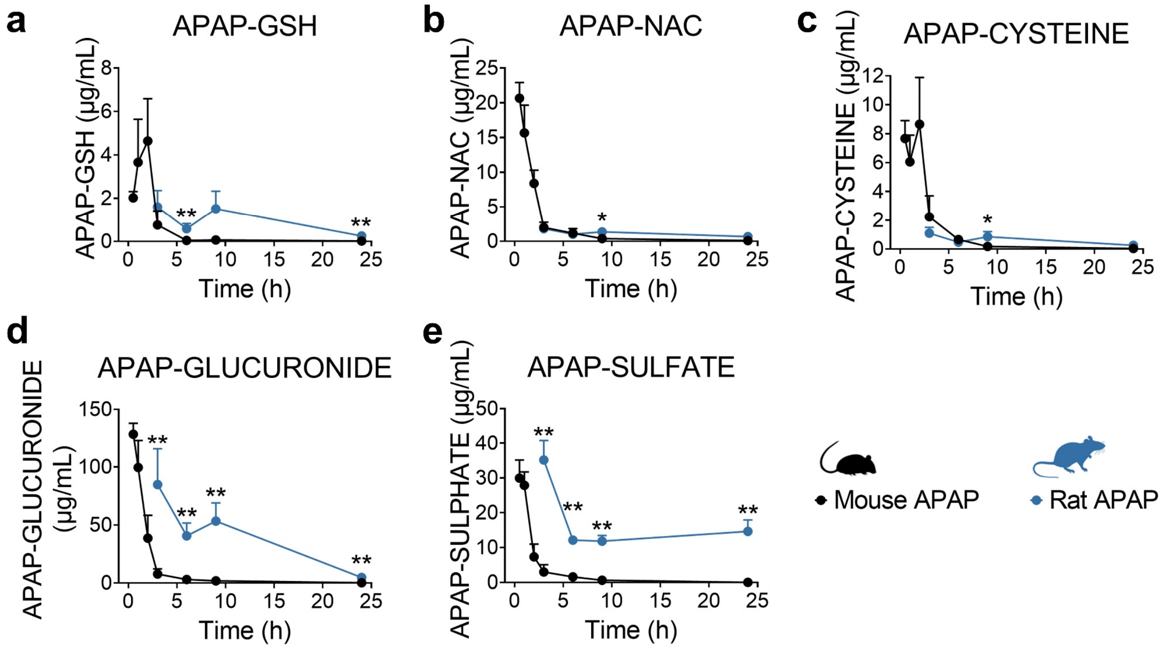


**Supplementary Figure 5.** Pharmacokinetic quantification of APAP metabolites (APAP-GSH (**a**), APAP-NAC (**b**), APAP-Cysteine (**c**), APAP-Glucuronide (**d**) and APAP-Sulfate (**e**)) in the plasma of C57Bl/6J mice (0.5, 1, 2, 3, 6, 9 and 24 h) and Sprague-Dawley rats (3, 6, 9 and 24 h) exposed to 300 mg/kg and 1000 mg/kg APAP, respectively. Values are mean ± SD (n=5). Statistical significance was determined between APAP-treated mice and rats where possible per time point (Mann-Whitney U test). P-values are denoted as *p<0.05, and **p<0.01.

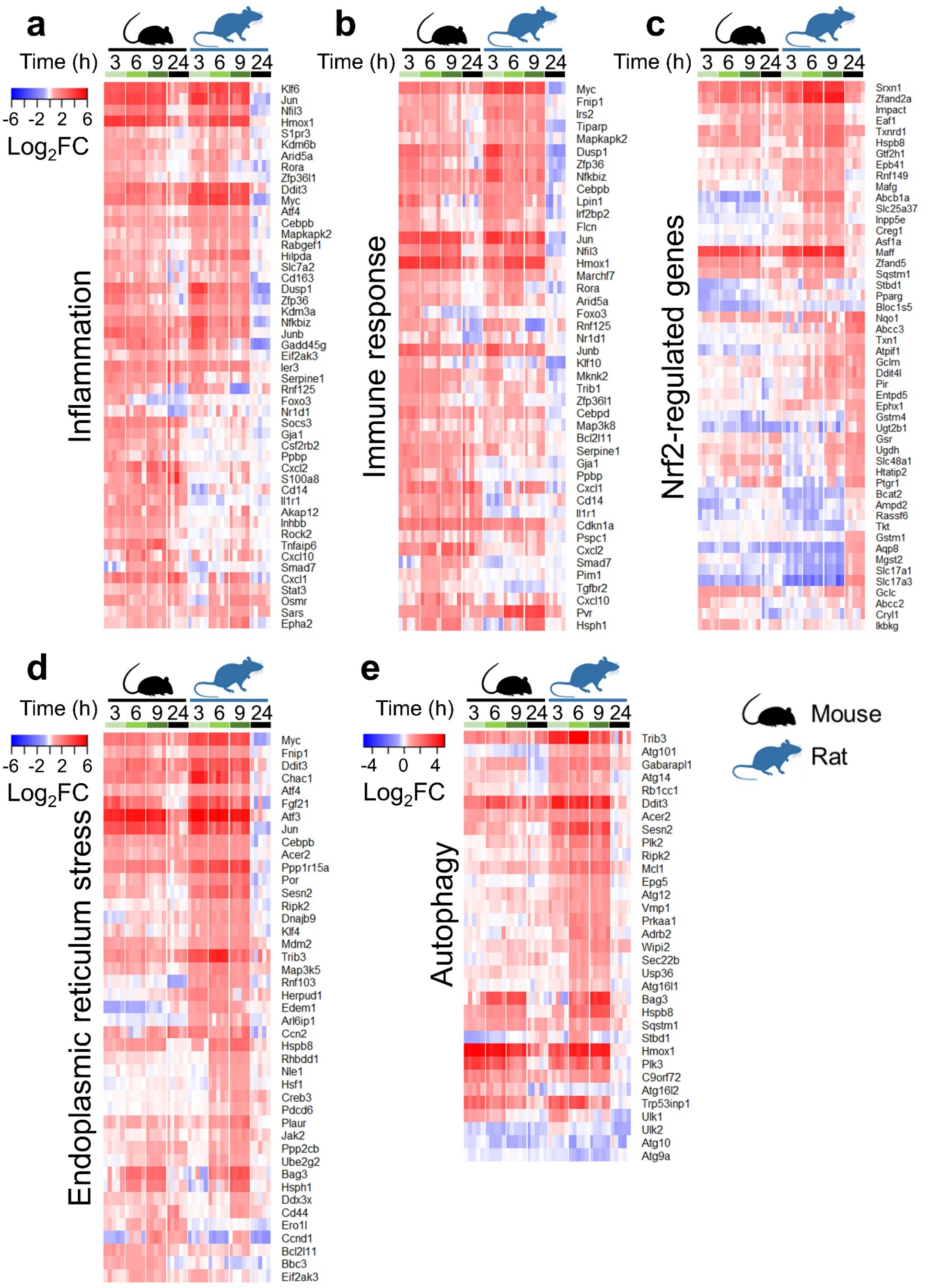


**Supplementary Figure 6.** Heatmaps showing changes in the expression of genes regulating (**a**) inflammatory, (**b**) immune, (**c**) oxidative stress, (**d**) endoplasmic reticulum (ER) stress responses, and (**e**) autophagy in the livers of C57Bl/6J mice and Sprague Dawley rats treated with 300 mg/kg and 1000 mg/kg APAP, respectively (n=5) at 3, 6, 9, and 24 hours. Data are expressed as $\log_2$ fold-change *versus* time-matched vehicle control animals.

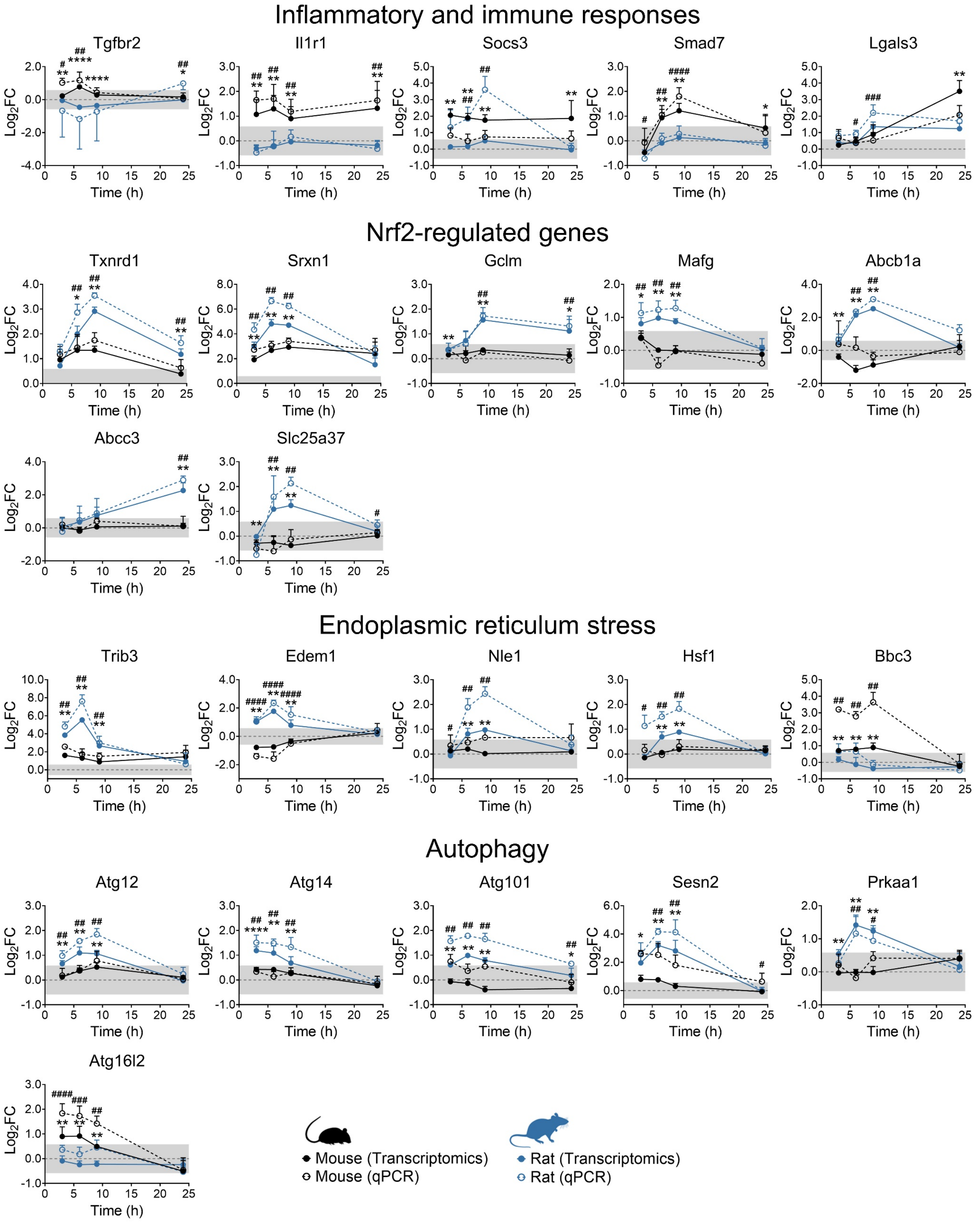


**Supplementary Figure 7.** mRNA expression levels of representative genes implicated in the indicated processes, as determined by qPCR analysis. Expression levels were normalised to GAPDH. Data are expressed as $\log_2$ fold-change versus time-matched vehicle control animals. Values are mean ± SD (n=5). Unpaired t-test or Mann-Whitney U test, as appropriate. P-values are denoted as *p<0.05, **p<0.01, or ****p<0.0001, comparison of mouse transcriptomics versus rat transcriptomics, or $^{\#}$p<0.05, $^{\#\#}$p<0.01, $^{\#\#\#}$p<0.001, $^{\#\#\#\#}$p<0.0001, comparison of mouse qPCR versus rat qPCR.

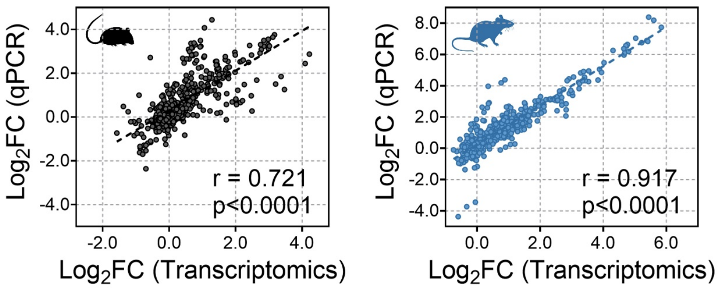


**Supplementary Figure 8.** Linear correlations between transcriptomics and qPCR data. Values are expressed as $log_2$ fold-change versus time-matched vehicle control animals. Pearson correlation coefficient (r) and p value for each species are shown in the plot.

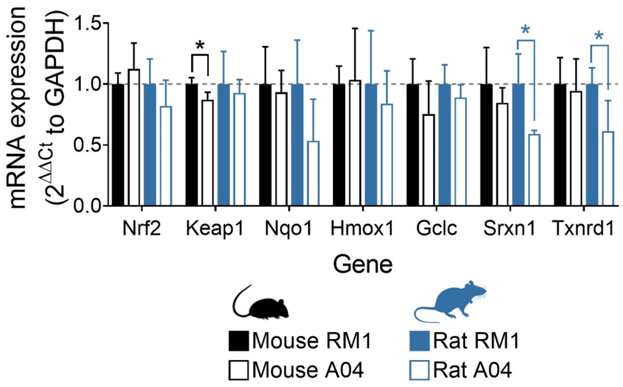


**Supplementary Figure 9.** mRNA expression levels of representative Nrf2-modulated genes in mice and rat fed with diets containing relatively high (A04, Safe, France) and low (RM1, Special Diet Services, UK) levels of arsenic. Male C57Bl/6J mice and male Sprague-Dawley rats were assigned to either RM1 or A04 diet for 7 days. Expression levels were normalised to GAPDH. Values are mean ± SD (n=4). Statistical significance was determined against RM1-fed animals (Unpaired t-test or Mann-Whitney U test, as appropriate). P-values are denoted as *p<0.05.

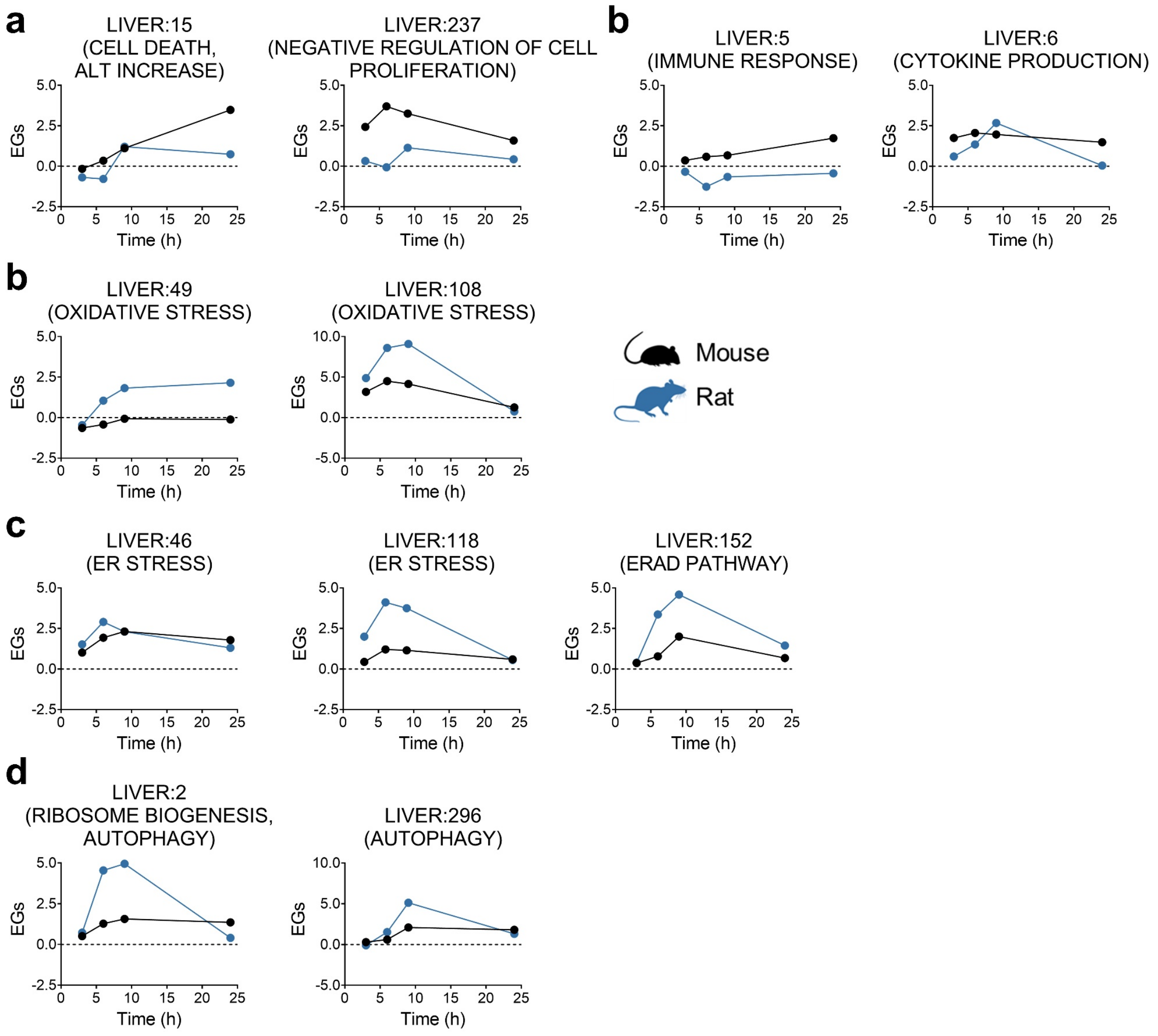


**Supplementary Figure 10.** Weighted gene co-expression network analysis (WGCNA) of differentially expressed genes in APAP-treated mice and rats using the TXG-MAPr web tool (https://txg-mapr.eu). A numeric Eigengene score (EGs) that aggregates fold-change values for the underlying genes in the module was calculated for both species at each time point. Relevant modules involved in the modulation of (**a**) cell death, (**b**) immune, inflammatory, (**c**) oxidative stress and (**d**) endoplasmic reticulum stress responses, and (**e**) autophagy**.** ALT, alanine aminotransferase; ER, endoplasmic reticulum; ERAD, endoplasmic-reticulum-associated protein degradation. See Supplementary File 3 for the full list of perturbed modules in both species.

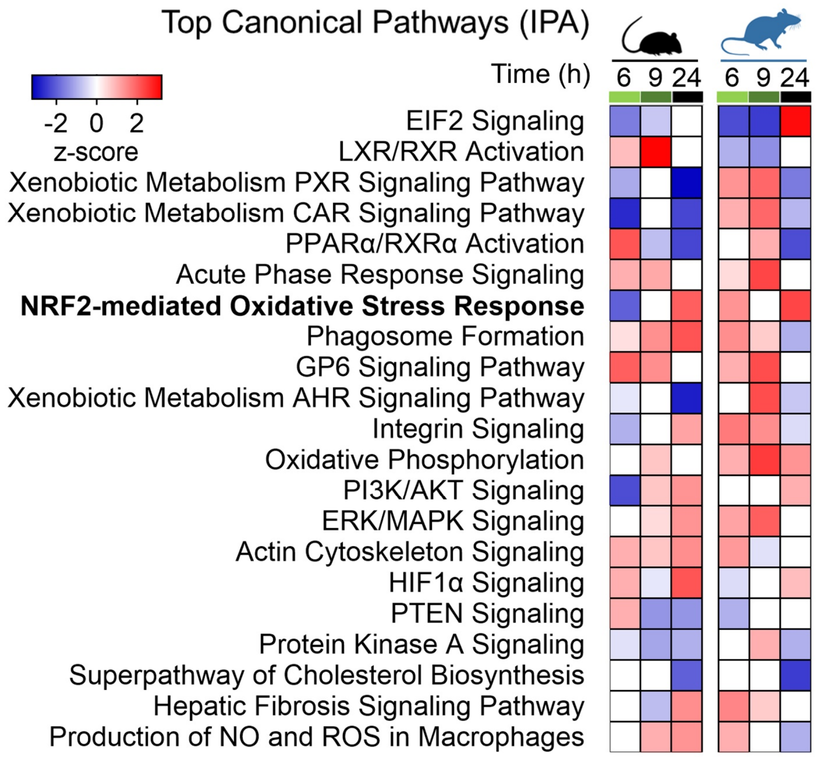


**Supplementary Figure 11.** Comparative analysis of top canonical pathways (IPA) in the liver of mice and rats treated with acetaminophen (APAP) 300 mg/kg and 1000 mg/kg, respectively (n=5) at 6, 9, and 24 hours (SWATH proteomics). For each function, a z-score was calculated at each time point against time-matched vehicle control animals.

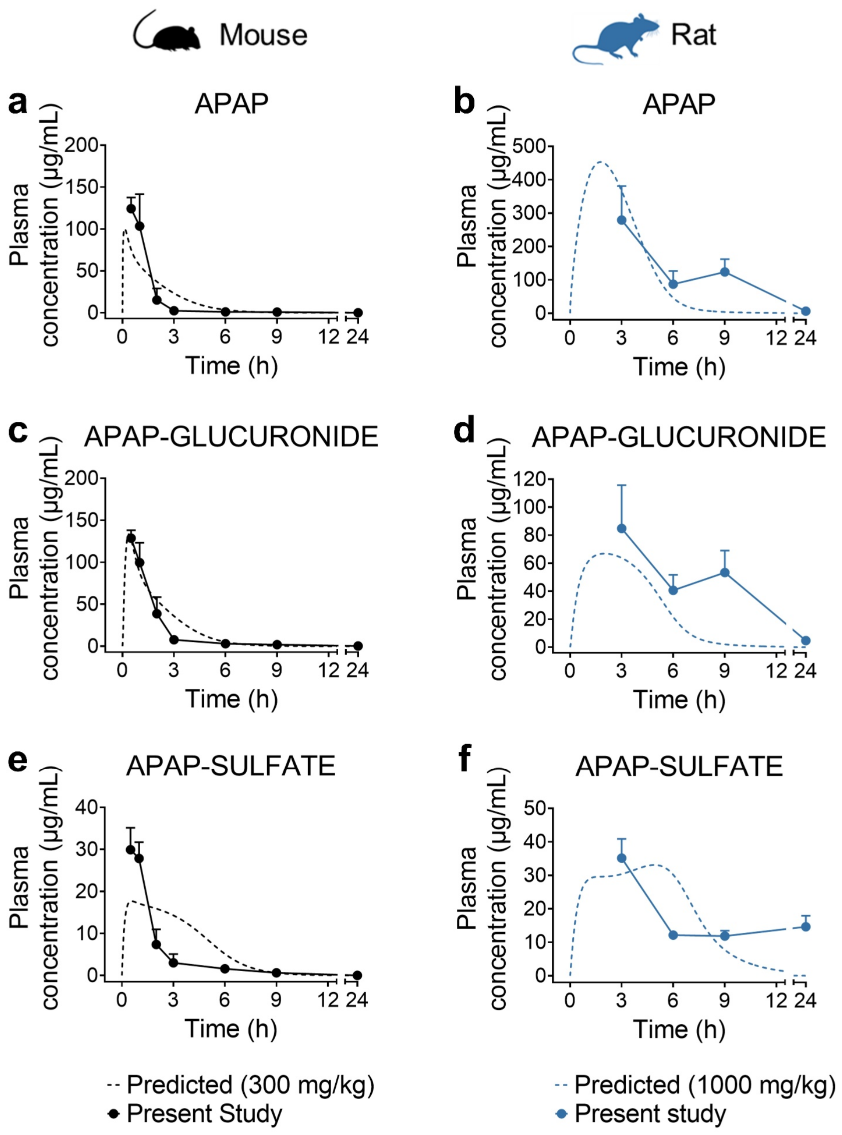


**Supplementary Figure 12.** Performance verification of physiologically-based pharmacokinetic (PBPK) models for simulation of APAP (**a-b**), glucuronidated metabolites (**c-d**), and sulfated metabolites (**e-f**). Results for the PBPK models (mouse on the left, rat on the right) show simulations (dotted lines) of a 300 mg/kg or 1000 mg/kg oral dose for mouse and rat, respectively, compared with the observed data from the in vivo study reported here (black or blue dots and solid lines; mean ± SD).

**SUPPLEMENTARY FILES**

**Supplementary File 1 – Calculation of hepatic NAPQI burden:** Total hepatic NAPQI burden based on glutathione depletion in mice and rats.

**Supplementary File 2 – GSEA:** Differentially expressed genes (DEGs) and Gene Set Enrichment Analysis (GSEA) in the livers of C57Bl/6J mice and Sprague Dawley rats treated with 300 mg/kg and 1000 mg/kg acetaminophen (APAP), respectively (n=5) at 3, 6, 9, and 24 hours. Selected GO processes shown in the main paper are highlighted in green.

**Supplementary File 3 – WGCNA:** Weighted gene co-expression network analysis (WGCNA) in the livers of C57Bl/6J mice and Sprague Dawley rats treated with 300 mg/kg and 1000 mg/kg acetaminophen (APAP), respectively (n=5) at 3, 6, 9, and 24 hours. The list of the top 50 most significant genes in each module (ordered by significance) and their relative $\log_2$ fold-change are also provided.

**Supplementary File 3 – Binned Protein Expression:** Basal protein expression levels grouped into bins. $\text{Log}_2$ transformed normalised protein expression values from SWATH proteomics in the livers of untreated (0 h) C57Bl/6J mice and Sprague Dawley rats (n=5) were ranked and grouped into 10 bins. Proteins with the lowest abundance were assigned to bin 1, and those with the highest abundance to bin 10. A bin value of 0 was assigned to proteins that were not detected.
